# Magnipore: Prediction of differential single nucleotide changes in the Oxford Nanopore Technologies sequencing signal of SARS-CoV-2 samples

**DOI:** 10.1101/2023.03.17.533105

**Authors:** Jannes Spangenberg, Christian Höner zu Siederdissen, Milena Žarković, Sandra Triebel, Ruben Rose, Christina Martínez Christophersen, Lea Paltzow, Mohsen M. Hegab, Anna Wansorra, Akash Srivastava, Andi Krumbholz, Manja Marz

**Affiliations:** RNA Bioinformatics and High-Throughput Analysis, Friedrich Schiller University Jena, Leutragraben 1, 07743 Jena, Germany; Institute for Infection Medicine, Christian-Albrechts-Universität zu Kiel and University Medical Center Schleswig-Holstein, Campus Kiel, Brunswiker Straße 4, 24105 Kiel, Germany; Labor Dr. Krause und Kollegen MVZ GmbH, Steenbeker Weg 23, 24106 Kiel, Germany; European Virus Bioinformatics Center 2, Leutragraben 1, 07743 Jena, Germany; FLI Leibniz Institute for Age Research, Beutenbergstraße 11, 07745 Jena, Germany

**Keywords:** Oxford Nanopore Technologies, raw ONT sequencing signal, differential RNA modifications, SARS-CoV-2, comparative analysis

## Abstract

Oxford Nanopore Technologies (ONT) allows direct sequencing of ribonucleic acids (RNA) and, in addition, detection of possible RNA modifications due to deviations from the expected ONT signal. The software available so far for this purpose can only detect a small number of modifications. Alternatively, two samples can be compared for different RNA modifications. We present `Magnipore`, a novel tool to search for significant signal shifts between samples of Oxford Nanopore data from similar or related species. `Magnipore` classifies them into mutations and potential modifications. We use `Magnipore` to compare SARS-CoV-2 samples. Included were representatives of the early 2020s Pango lineages (n=6), samples from Pango lineages B.1.1.7 (n=2, Alpha), B.1.617.2 (n=1, Delta), and B.1.529 (n=7, Omicron). `Magnipore` utilizes position-wise Gaussian distribution models and a comprehensible significance threshold to find differential signals. In the case of Alpha and Delta, `Magnipore` identifies 55 detected mutations and 15 sites that hint at differential modifications. We predicted potential virus-variant and variant-group-specific differential modifications. `Magnipore` contributes to advancing RNA modification analysis in the context of viruses and virus variants.

## Introduction

Oxford Nanopore Technologies (ONT) provides the possibility to sequence desoxyribonucleic acid (DNA) and ribonucleic acid (RNA) directly without any amplification, which would erase nucleotide modifications. ONT uses flow cells containing nanopores with sequencing sensors. Nucleotides are pulled through nanopores. The pore is integrated into a membrane to which a voltage is applied. The measured electrical current within the nanopore is characteristic of the nucleotides within the pore. Five nucleotides are measured at a time in the most narrow part of the pore. The change in the electric current enables the basecalling procedure, which can be identified by deep learning or statistical models [1, 2, 3, 4].

Severe acute respiratory syndrome coronavirus type 2 (SARS-CoV-2) is a single positive-stranded RNA virus with a genome size of about 30kb and a 3’ poly-A tail [5, 6]. Since the end of 2019, various SARS-CoV-2 variants have emerged that differ from the original Wuhan variant by defined genomic mutations and resulting amino acid substitutions, particularly in the spike protein [7]. Bioinformatics pipelines like poreCov [8] and others [9, 10] contribute significantly to learning more about the virus and containing the COVID-19 pandemic by identifying and monitoring emerging variants. They also provide insights into potential RNA secondary structures [11, 12].

In the mammalian cell, RNA modifications influence, among other things, the formation of secondary and tertiary structures of transfer RNAs and the translation, stability, and localization of messenger RNAs (mRNAs) [13]. Previous studies have shown that positive single-stranded RNA viruses like *Picornaviruses* or *Flaviviruses* contain RNA modifications that play a significant role in viral infection and replication and even influence the host antiviral innate immunity [14, 15, 16, 17]. The replication of various coronaviruses also appears to be affected by RNA modifications [18]. The growing understanding of the importance of such modifications is accompanied by increased activities to detect them by direct RNA sequencing (DRS) [19, 20, 21, 22]. In ONT sequencing, modifications, as well as mutations, lead to shifts of the ONT signal [23, 24] and changes in the translocation speed [24].

Several machine-learning approaches have been developed to detect specific modifications. For N^6^-Methyladenosine (m6A), methods employed include support-vector machines as in `EpiNano` [21], neural networks like `m6Anet` [19], as well as random forests in `MINES` [22]. However, due to the lack of training data, the individual ONT signals cannot be reliably assigned to one of the many modifications. So far, there are only approaches that predict a modification at a nucleotide position with a certain probability.

Another strategy is a generic detection of modification sites without assignment to a specific modification by comparing two ONT samples. Some tools, e.g., `DRUMMER` [25], or `ELIGOS` [26], focus on different patterns of sequencing errors after basecalling, introduced by RNA modifications.

Other tools, such as `Nanocompore` [20], `Yanocomp` [27], or `xPore` [28], analyze the raw ONT signal directly. These tools compare the raw ONT data of two samples from a control and a wild-type condition to find differentiating signals [20, 27, 28]. The control sample is thereby depleted of modifications, and the wild type should carry modifications. All three tools are limited to comparing samples of the same species and classifying differentiating signals as modifications, neglecting mutations.

Here, we present `Magnipore`, a Python 3 pipeline for comparative ONT data analysis on signal level, Fig. 1. `Magnipore` takes two reference `FASTA` files to compare two distantly related genomes with alignable homologous regions. The program creates signal distribution models from the raw signal in the `FAST5` files. These distributions are compared between alignable regions. It predicts global differential signals between two samples. `Magnipore` classifies those into mutations and modifications using reference sequences detecting epitranscriptomic and transcriptomic differences. For signal segmentation and resquiggling, `Magnipore` uses `nanopolish` [29], which requires basecalled reads. The basecalls can be produced using `Guppy` within the pipeline, or the user can provide them. We map the basecalled reads with `minimap2` [30] to segment the raw ONT signals corresponding to the bases in the reference sequence using `nanopolish eventalign`. For each base in the reference, we calculate a Gaussian signal distribution from all reads mapped to this base.

**Figure 1:**
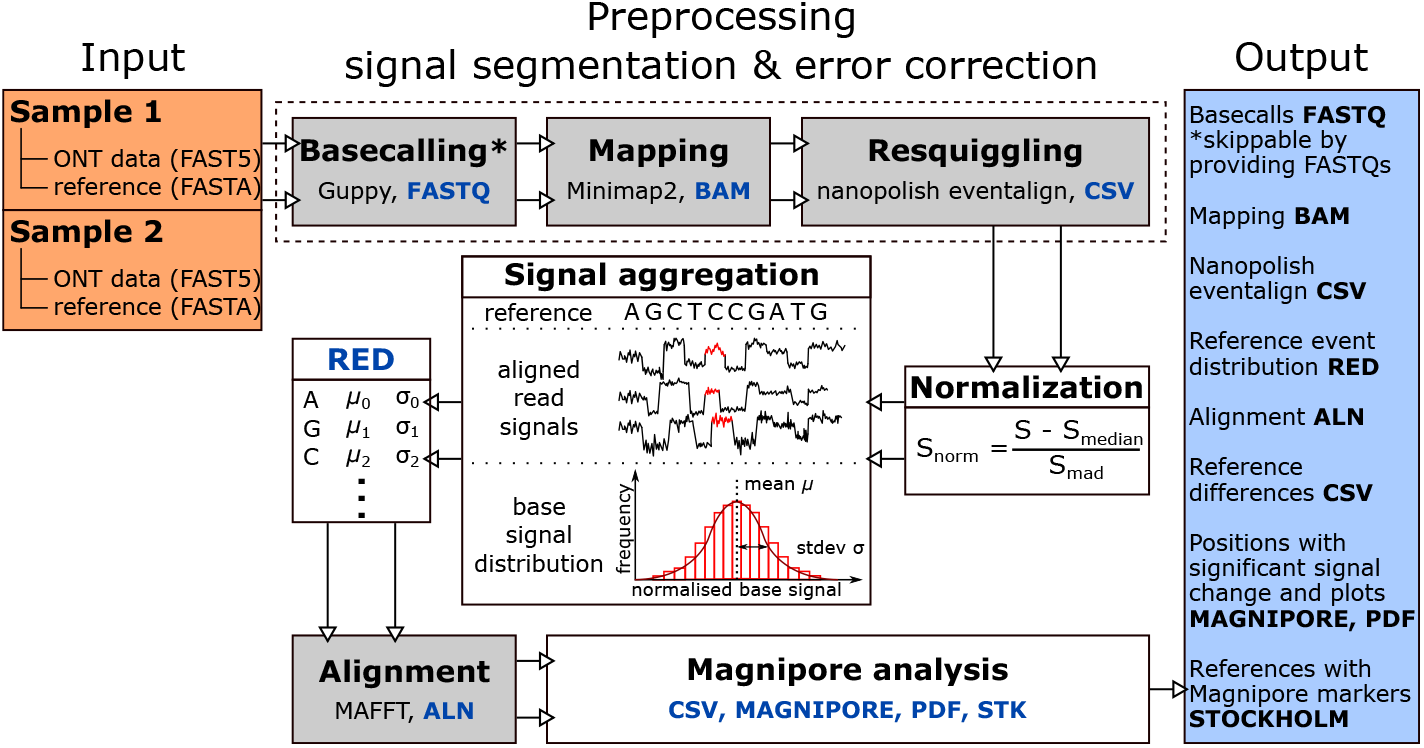
Workflow of `Magnipore`. A full `Magnipore` run starts with the preparation of two input samples. The incoming sample can be basecalled using `Guppy`. The `FASTQ` files are aligned to a given reference sequence using `minimap2`. The resulting `BAM`, `FASTQ`, and `FASTA` references are provided to `nanopolish eventalign` to resquiggle and segment the raw ONT data. The segmented read signals are then normalized. For each input sample and for each reference position, the segmented base signals from `nanopolish` are used to approximate a Gaussian signal distribution. To know which positions need to be compared, the input references will be aligned using `mafft`. If both samples share the same reference, then it must be provided in both inputs. The Gaussian signal distributions are compared for each aligned position of both samples. Unaligned positions with gaps are written to the `INDEL` file. Positions of significance surpass a significance threshold calculated according to Eqn. 1, which we will call sites. These significant sites can be ordered and filtered according to their threshold distance (TD) score, coverage, and other output values in the `Magnipore` file, which aids downstream analysis. Significant sites are also marked next to the provided reference sequences in a `STOCKHOLM` file for further analysis. Plots, as in Fig. 5, are also created.

As a study case, we aim to identify variant-specific mutations and modifications of SARS-CoV-2 using `Magnipore`.

## Materials and Methods

### Preparation of total RNA

Nucleic acids from nasopharyngeal swabs were analyzed for the presence of SARS-CoV-2 genome equivalents using a triplex real-time RT-PCR [31]. If the sample tested positive for SARS-CoV-2, the underlying strain was determined by whole genome sequencing. For this purpose, the NEBNext® ARTIC SARS-CoV-2 Companion Kit (New England Biolabs, Ipswich, MA, USA) and a MinION system (Oxford Nanopore Technologies plc., Oxford Science Park, UK) were used according to the manufacturer’s instructions. The principle of this procedure is to reverse transcribe the viral RNA into complementary desoxyribonucleic acid (cDNA) and amplify it with a mixture of primers so that the overlapping PCR fragments cover the entire viral genome. These are purified, barcoded, and sequenced. The primers are regularly adapted to newly emerging variants. Data were analyzed using the poreCov workflow and assigned to a Pango lineage [8, 32]. Aliquots of SARS-CoV-2-containing nasopharyngeal swab specimens in phosphate-buffered saline were then shaken in cell culture medium and placed in 48-well plates seeded with Vero cells, which were then incubated under standard conditions at 37°C for several days. At least one to two more passages on Vero cells followed after that. Then an aliquot of the supernatant was tested for the presence of SARS-CoV-2 RNA [31]. If SARS-CoV-2 genome equivalences were detectable, Vero cells seeded in T25 flasks were inoculated. After several days RNA was prepared using the RNeasy kit (Qiagen, Hilden, Germany) according to the manufacturer’s instructions. All virus cultivation and RNA extraction steps were performed under biosafety level-3 conditions. The resulting SARS-CoV-2 isolates were also sequenced according to the ARTIC protocol and used for neutralization experiments or other purposes [32]. The sequencing of the original samples was carried out for epidemiological reasons or according to the requirements of the Coronavirus Surveillance Ordinance (CorSurV) of the Federal Republic of Germany. The provision and sequencing of viral RNA in Jena are covered by a positive vote of the Ethics Committee of the Medical Faculty of Kiel University (D467/20).

### Direct RNA sequencing

The library was constructed using ONT protocol with the direct RNA sequencing and SQK-RNA002 kit. The manufacturer’s instructions were followed with minor changes listed to achieve the suggested recovery aim of about 200 ng of the final library and to obtain longer reads: 1. Before library preparation, the sample was cleaned, concentrated, and size selected for longer reads with RNAClean XP beads, with a ratio of up to 4:3 (sample:beads); 2. The starting sample amount was increased from 1 μg to 4 μg RNA; 3. The amount of the RNA Control Strand (RCS) was used for samples 5 and 9 (see Tab. 1 as dilution 1:10, and for other samples not added at all; 4. As we noticed a lot of free adapters, RNA Adapter Mix (RMX), we reduced the amount from 6 μl to 3 μl; 5. For the Omicron BE.1 and BF.1 samples, incubation times were prolonged so that adapters had more time to ligate. The reaction with the RT adapter (RTA) and RMX time was prolonged to 15 min and 20 min, respectively. The library was loaded onto R9.4.1 flow cell and sequenced on a MinION Mk1B device.

**Table 1:** SARS-CoV-2 RNAs included in this study, for details see STab. S1. Superscript ^a, b^, and ^c^ denote different patient isolates of the same Pango lineage. All samples were sequenced using an ONT MinION Mk1B platform with an R9.4.1 flow cell. *The raw ONT data and basecalled `FASTQ`s can be found in the SRA BioProject: PRJNA907180. **All reference sequences are available via the OSF database https://osf.io/evc6k/. Samples were sequenced using direct RNA sequencing (DRS) on an ONT MinION platform. We show results from `Magnipore` comparisons using B.1.1.7^a^ as the reference sample.

| Pangolin | Nextclade (WHO) | Sample* | Reference** | Gisaid ID |
| --- | --- | --- | --- | --- |
| B.1.513 | 20C | MinION DRS | MinION ARTIC | — |
| B.1<sup>a</sup> | 20C | MinION DRS | MinION ARTIC | — |
| B.1<sup>b</sup> | 20C | MinION DRS | MinION ARTIC | — |
| B.1<sup>c</sup> | 20C | MinION DRS | MinION ARTIC | — |
| B.1.177.86 | 20E | MinION DRS | MinION ARTIC | — |
| B.1.1 | 20B | MinION DRS | MinION ARTIC | — |
| B.1.1.7<sup>a</sup> | 20I (Alpha) | MinION DRS | MiSeq ARTIC | EPI_ISL_862139 |
| B.1.1.7<sup>b</sup> | 20I (Alpha) | MinION DRS | MiSeq ARTIC | EPI_ISL_862140 |
| B.1.617.2 | 21J (Delta) | MinION DRS | MiSeq ARTIC | EPI_ISL_2500366 |
| BA.1.17.2 | 21K (Omicron) | MinION DRS | MiSeq ARTIC | EPI_ISL_7929223 |
| BA.1.18 | 21K (Omicron) | MinION DRS | MiSeq ARTIC | — |
| BA.2.9 | 21L (Omicron) | MinION DRS | MiSeq ARTIC | EPI_ISL_9553935 |
| XBB.1.5 | 22F (Omicron) | MinION DRS | MinION ARTIC | — |
| BA.4.1 | 22A (Omicron) | MinION DRS | MinION ARTIC | — |
| BE.1 | 22B (Omicron) | MinION DRS | MinION ARTIC | — |
| BF.1 | 22B (Omicron) | MinION DRS | MinION ARTIC | — |

### Magnipore requirements

We recommend using a conda environment for `Magnipore`. Required packages and tools except `Guppy` can easily be installed with conda via https://anaconda.org/JannesSP/magnipore or with the YAML file in the GitHub repository https://github.com/JannesSP/magnipore. A detailed list of the packages, tools, and their versions can be found in Tab. 2.

**Table 2:**
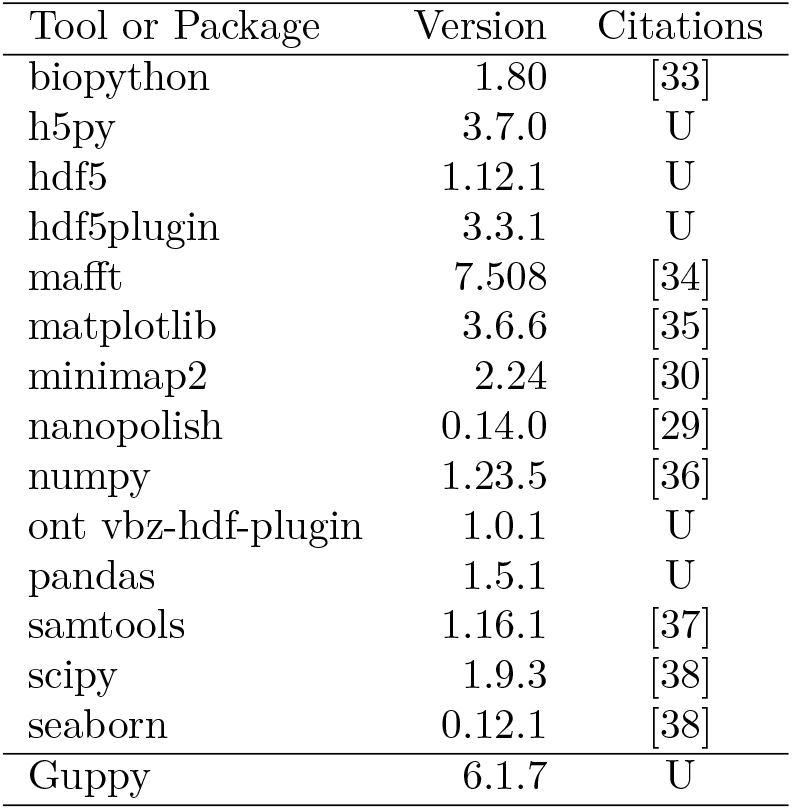
Tools included in this study. To run the `Magnipore` pipeline, different external tools are necessary. `Guppy` must be installed and downloaded from nanoporetech, which is only available to customers. All other tools can be installed with conda. `Magnipore` was tested with all listed external tools and their corresponding version. Some tools or packages are not published (U – unpublished). This is the case for h5py (https://zenodo.org/record/6575970) , hdf5 (https://www.hdfgroup.org/HDF5/), hdf5plugin (https://github.com/silx-kit/hdf5plugin) , ont vbz-hdf-plugin (https://github.com/nanoporetech/vbz-h5py-plugin) , pandas (https://zenodo.org/record/7658911) and Guppy (https://nanoporetech.com/community).

### Magnipore input

`Magnipore` requires two multi-`FAST5` files and two reference `FASTA` files as input for both samples (Fig. 1). In case the samples originate from the same biological sample for two different environmental conditions, the same reference file must be provided for both sample inputs. `Magnipore` requires basecalls for `nanopolish`’s signal segmentation. The user provides either those basecalls with --path_to_first_basecalls PATH and --path_to_sec_basecalls PATH or `Magnipore` needs the `Guppy` binary and model path with `--guppy_bin PATH` and `--guppy_model PATH` to basecall the `FAST5` data.

### Basecalling with Guppy

Guppy v6.1.7 is available for ONT customers and can be obtained via their community site https://community.nanoporetech.com. `Magnipore` uses Guppy with the parameters `--disable_qscore_filtering --calib_detect -c guppy_model` if no `FASTQ` file is provided.

### Mapping with minimap2

In the next step, `Minimap2 v2.24` [30], using the parameters `-a -x splice -k 14` maps the reads to the provided reference to run `nanopolish eventalign` v0.14.0 [29]. For that we executed `nanopolish` index with the parameters to execute `nanopolish eventalign` with `--summary=summary.csv --scale-events --signal-index -t threads`.

### Resquiggling with nanopolish

`Magnipore` needs the ONT signal segment assignment for each base to calculate a normalized signal distribution for each position. `nanopolish eventalign` resquiggles and segments the raw ONT signal from the `FAST5` files using the mapping from `minimap2` [29]. `nanopolish eventalign` can only process those reads that could be mapped to the reference sequence. Resquiggling the reads means that `nanopolish eventalign` corrects the bases in the basecalled reads according to the given reference sequence, integrated k-mer models from ONT, and the raw ONT signal. `nanopolish eventalign` can not distinguish between basecalling errors or real mutations. It assumes that the reference sequence is the ground truth and will correct every base from the reads using the corresponding region in the reference sequence. Additionally to the resquiggling, `nanopolish eventalign` assigns signal segments to each reference position.

### Normalization

The pico Ampere (pA) signal for each read is normalized by the subtraction of the median pA value from all measurements and scaled by its pA median absolute deviation. The normalization removes measurement biases between sequencing runs, pores, and sensors in a single sequencing run. After normalizing the raw ONT signal, we can compare the measurements across different reads and sequencing runs.

### Signal aggregation per position

Then, `Magnipore` aggregates the ONT signal for each reference position from the segmented reads provided by `nanopolish eventalign` and calculates the mean and standard deviation. We employ a Gaussian distribution for each position to approximate the signal distribution from all reads in the signal aggregation step (Fig. 1). This provides a good approximation of the ONT signal distribution, as most resemble a Gaussian distribution (SFig. S4). The mean and standard deviation are stored in the reference event distribution `RED` file, containing the reference position, base, motif, signal mean, and standard deviation. The file also contains quality values such as the coverage, distribution density, how many data points are within the 99th percentile of the Gaussian distribution, and more for downstream analysis. `Magnipore` reuses this file by default for comparisons with other samples to save calculation time by default.

### Magnipore: Position-wise signal comparison between samples

If the `RED` files for two samples are created, mafft v7.508 [34] auto alignment mode (`--auto`) is used to align the sample references. The mapping is used to correctly compare the signals per positions of two different individuals positionwise. The signal distributions for the alignment of position *i* from the reference of sample 1 and position *j* from the reference of sample 2 are considered to be of significance if the absolute difference of their means is larger than the average of their standard deviations, that is:

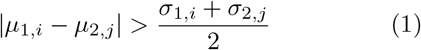

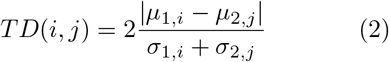

Alignment positions are called sites. Furthermore, the threshold distance (TD) score provides a convenient order for the significant sites. The significant sites are given by *TD*(*·,·*) ≥ 1, and the insignificant sites are given by *TD*(*·,·*) *<* 1. A significant signal change between position *i* of one sample and *j* of the other sample can be caused by molecular changes between both in close proximity to the investigated site. A molecular change can be a mutation or a modification. `Magnipore` classified a significant signal change into mutation or potential modification sites using the provided reference sequences. The classification of significant signals is further described in Tab. 3. `Magnipore` is available at https://github.com/JannesSP/magnipore.

**Table 3:**
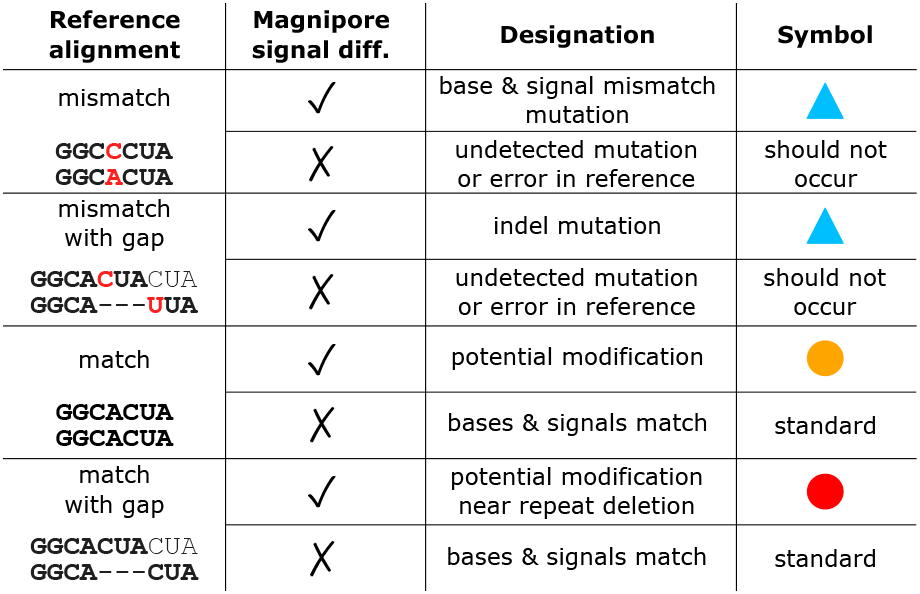
Biological interpretation of `Magnipore` events. In all cases, base A in the middle of the 7-mers in bold letters is the base of interest with a significant signal change.

### Correlation with RNA secondary structures

To investigate the correlation between potential modification sites and possible RNA secondary structures, we used an alignment-based approach: We clustered complete genomes of the genus *Betacoronavirus* (taken from NCBI virus [39], submitted before June 2020) with `MMSeqs2` [40] and identified for each cluster a centroid sequence as representative. We extracted a fragment of 100 *nt* around each significant `Magnipore` site (>40 %) derived from the genome BA.2.9 eligible for a modification. Homologous regions were used to construct a sequence-secondary-structure alignment by `mLocARNA` [41]. `VARNA` [42] and RNAalifold [43] were applied for visualization with the following parameters: `--aln --noLP --noPS --mis -r -d2 --cfactor 0.6 --nfactor 0.5`. We calculated the p-value of each prediction based on a dinucleotide shuffling implemented in a Python script available on GitHub^1^.

## Results and Discussion

### Magnipore classifies significant sites into mutation and potential modification

`Magnipore` classifies these significant sites into mutation and potential modification sites. It ignores the gaps in the reference alignment for the classifications and only uses sequence similarity. `Magnipore` compares the 7-mer sequence context across gaps.

Significant differential ONT signals (sites) found by `Magnipore` can appear in different types of events (Tab. 3): The left column in Tab. 3 depicts a comparison of the two input sequences in an alignment.

In case the two reference sequences have a mismatch with or without a gap and `Magnipore` identifies an expected signal change (√), then we interpret this to be a mutation site (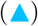), whereas no identified signal change should not occur.

If the sequences are identical (match) but `Magnipore` identifies a signal change, we assume this to be a potential modification (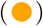), whereas no signal change is the standard case.

In case of a gap resulting in two possible alignments of equal value, `Magnipore` classifies a significant signal change as a potential modification site (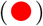). In both possible alignments, the sequence context is the same.

These cases can be further investigated in the output of `Magnipore`. `Magnipore` produces a `STOCKHOLM` file for each comparison in which all sites with a significant signal change are marked with an X in the sample alignment, see Fig. 2.

**Figure 2:**
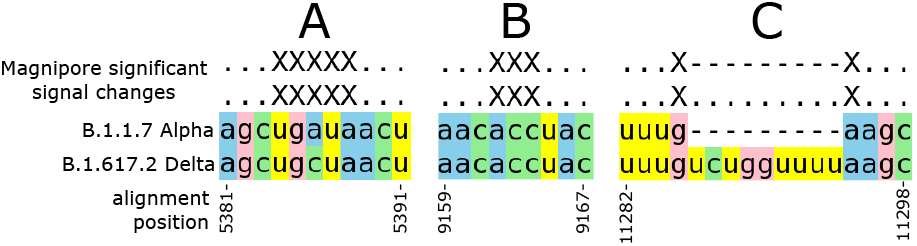
`Magnipore` provides the reference sequence with marked significant sites as a `STOCKHOLM` file. This file stores an alignment of marked sites with the references to visualize the context of the significant sites for further investigations. **A** shows a marked region that results from a substitution of the A in the Alpha variant to a C in Delta. **B** is an example for a site with no mutational context in the Illumina reference sequence from GISAID, that was created from the same original sample. **C** visualizes that also insertion and deletion regions result in significant sites.

We present `Magnipore` using the example comparison of B.1.1.7^a^ Alpha and B.1.617.2 Delta as they were the two first variants of concern in our datasets. Variants of concern may exhibit increased transmissibility and/or escape antibody-mediated neutralization [44]. These samples are therefore very interesting for research.

### Magnipore detects 89.1% of all mutations in the SARS-CoV-2 pairwise sample comparisons

To determine the performance of `Magnipore` to detect differential ONT signals, we analyze how many mutations `Magnipore` can identify in pairwise SARS-CoV-2 comparisons. `Magnipore` with default parameters and no coverage filter can detect 89.1% of all mutations within all our SARS-CoV-2 comparisons. A mutation is a difference in the alignment of the reference sequences between two samples, including substitutions, insertions, and deletions (see Tab. 3, blue triangle). We exclude those differences where at least one of the reference sequences shows the character N instead of A, C, G, or T/U. A mutation is captured if `Magnipore` reports at least one significant site in close proximity to three bases upstream or downstream. Across 120 pairwise comparisons 13 978 non N mutations are present in the reference sequences. `Magnipore` identifies 12 460 of those using the significance threshold Eqn. 1 with Gaussian distribution models. The 12 460 identified mutations out of 13 978 form a ratio of 89.14%.

Looking at the pairwise comparisons individually, the median of detected non N mutations is 89.1% while the mean is 87.2%, Fig.4. We calculated the fraction of found non N mutations for each comparison first, then we averaged over all fractions. The lowest fraction of detected mutations is 50% in the comparison of B.1^a^ against B.1^b^ where `Magnipore` finds 2 out of 4 mutations. The highest is 100% in B.1^a^ compared to B.1^c^ with 3 mutations.

In the pairwise `Magnipore` comparisons of B.1.1.7^a^ Alpha against all other samples we filtered for a coverage of 10 and higher, Fig. 3. In the B.1.1.7^a^ Alpha comparisons the mean fraction of detected mutations is 80.07%. It ranges from 51.0%, B.1.1.7^a^ compared to XBB.1.5 Omicron, to 91.9%, B.1.1.7^a^ compared to B.1.513.

**Figure 3:**
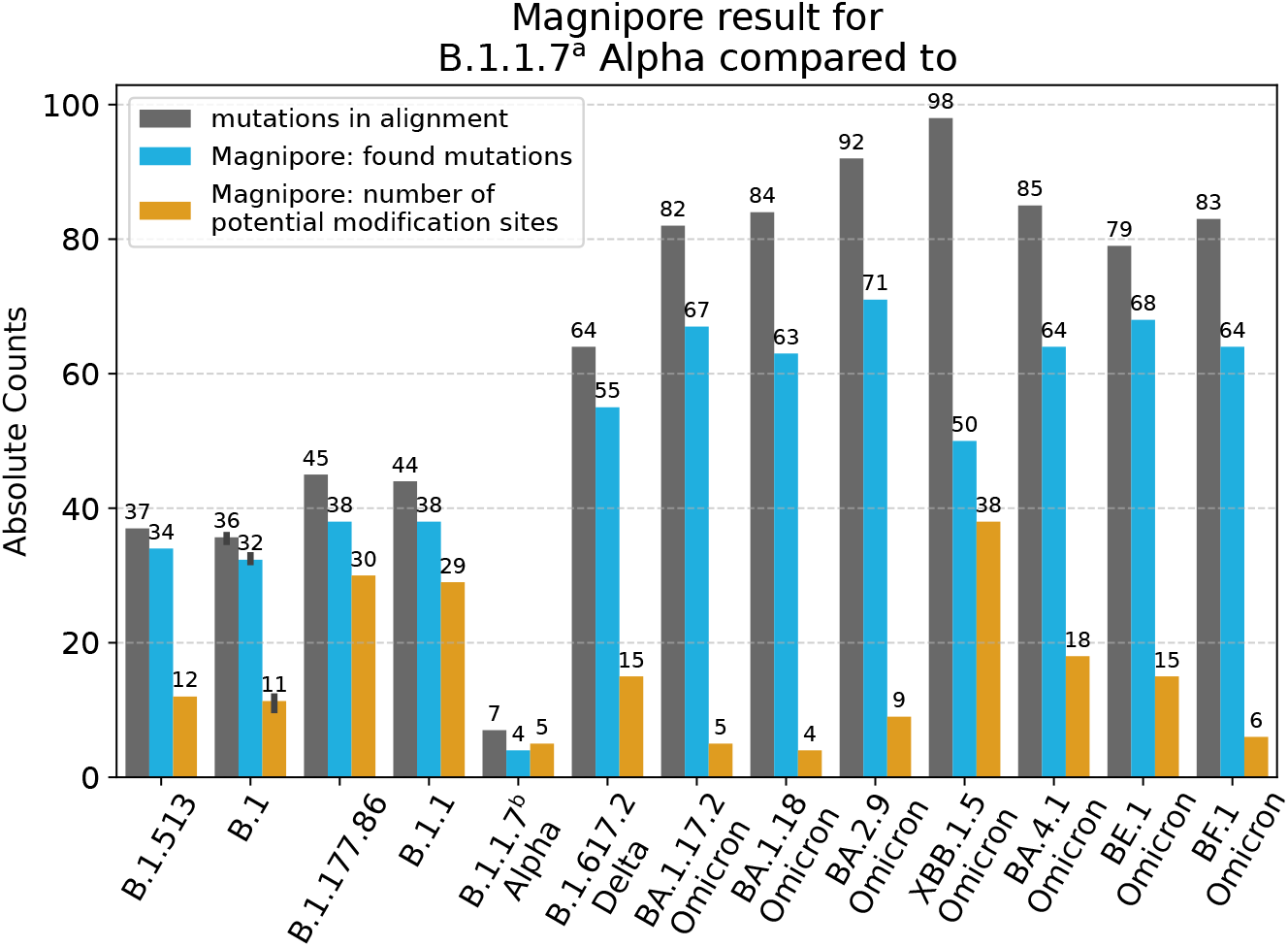
Results by `Magnipore` for B.1.1.7^a^ Alpha in comparison with all other SARS-CoV-2 strains from Tab. 1. For the comparisons we used a coverage filter of 10. Sites with a coverage below 10 were discarded. We counted the number of mutations between the reference sequences – shown in grey; blue – fraction of identified mutations with `Magnipore`; orange – potential modifications. In case of B.1, we had three samples. The standard deviation of the counts is indicated by the black dash on top of the bars.

**Figure 4:**
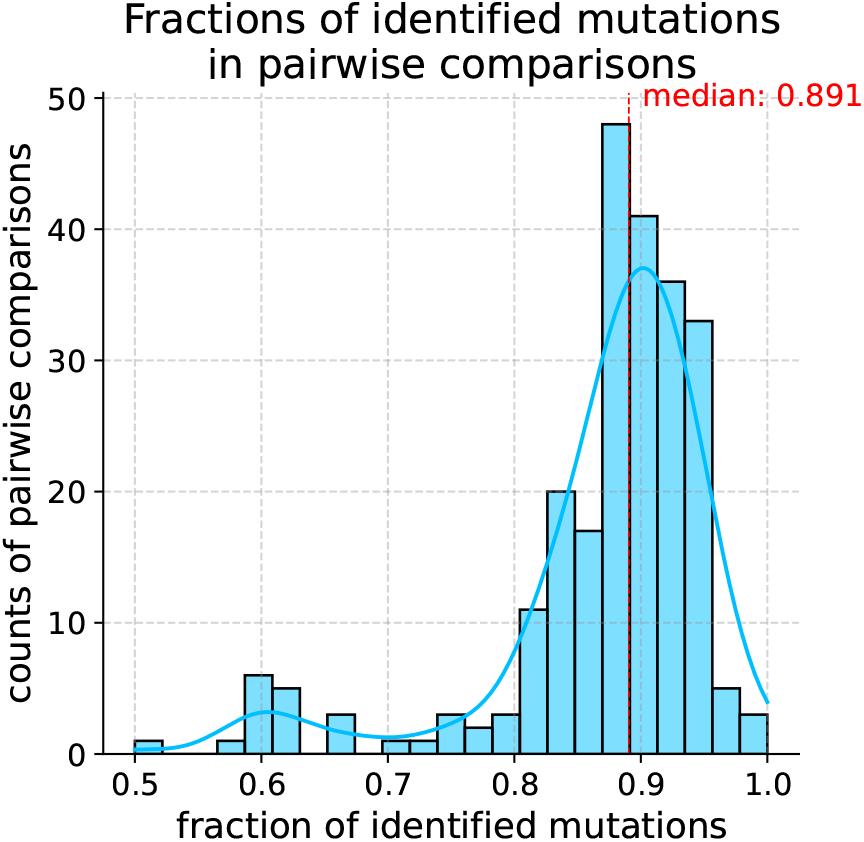
Distribution of fractions of identified mutations (detected non N mutations divided by non N mutations in the references) for all pairwise comparisons without filtering for the coverage. The median number of identified mutations is 0.891, the mean is 0.872.

`Magnipore` is not able to detect 10.9% of the present mutations. We assume that there are three reasons why `Magnipore` misses 10.9 % of mutations: 1. Inaccurate signal segmentation and resquiggling; 2. Gaussian distributions to capture all types of signals are suboptimal; and 3. Our significance threshold to detect differential signal distributions is not optimal or the signal distribution of the mutated bases are too similar. In both cases they do not show a significant differential signal.

### Inaccurate signal segmentation and resquiggling

`Magnipore` uses `nanopolish eventalign` to resquiggle and segment the raw ONT signal.

The segments should correspond to a measured sequence of bases, and the segment borders resemble the signal transition of bases measured in the nanopore. Additionally, `nanopolish eventalign` uses the given reference sequence to correct errors within the basecalled reads. This way, bases are associated with the signal segments. Thereby, `nanopolish` erases true mutations and falsely corrects them using the reference bases. It is a false error correction, as haplotypes are erased, and wrong bases are associated with signal segments. That can lead to skewed or mixed signal distributions for a single reference position, SFig. S4. Another reason for mixed distributions is inaccurate signal segmentation, which can happen if the base transmission border is misplaced.

### Gaussian distributions to capture all types of signals are suboptimal

We use Gaussian distribution models to approximate each sample’s true signal distribution per base. In many cases, the Gaussian distribution model is a reasonable approximation. In other cases, the true signal distribution does not resemble a Gaussian distribution, SFig. S4. The signal distribution shows a mixture of multiple Gaussian distributions in these cases. Such behavior can be caused by inaccurate segmentation or false error correction from the previous case. Our current model cannot capture this behavior. These signal distributions skew the distribution model, which might be why some mutations remain undetected.

### Significance threshold not optimal or mutated signals not differential

Another reason for undetected mutations could be that Eqn. 1 is a suboptimal significance threshold to detect all mutations. The signal distributions of mutated signals can sometimes be too similar and therefore do not show a differential signal. One example is the 5-mer models of AATCA and AAGCA that are provided by ONT and used by `nanopolish`. The k-mer distribution models can be found in [nanoporetech’s github](https://github.com/nanoporetech/kmer_models/blob/master/r9.4_180mv_70bps_5mer_RNA/template_median69pA.model). The 5-mer models of AATCA and AAGCA have very similar distributions, although they have different bases in the middle of the 5-mer. Their mean values are too similar and would not be detected with Eqn. 1. We also tried to use the Kullback-Leibler (KL) divergence with an empirical threshold to find differential signals, SFig. S1.

### KL divergence as an alternative significance threshold

The KL divergence is a well-known statistical method to describe diverging or overlapping distributions [45]. A small KL divergence value means less diverging distributions, while a large value means strongly diverging distributions. We provide the KL divergence value in the `Magnipore` output and compare it to then TD score. After looking at the KL divergence distribution in the comparison of B.1.1.7^a^ Alpha and B.1.617.2 Delta, SFig. S1, we decided to set the cutoff for a significant signal change to 1. Each site with a KL divergence above or equal to 1 is significant. We have no gold standard or ground truth data for our samples, so we evaluated the performance of `Magnipore` according to the known mutations between the samples. There are 64 non N mutations between the B.1.1.7^a^ and B.1.617.2 samples, Fig. 3. Using the TD score and a coverage filter of 10, we found 146 mutation signals clustering around 55 non N mutations. Using the KL divergence threshold of 1 and a coverage filter of 10, we found 117 mutation signals clustering around 53 non N mutations. Both results share 114 mutation signals, which means with the TD score, we can detect more mutation signals and more identified mutations, SFig. S2, S3. Using the TD score we do not need to estimate a significance cutoff for every sample comparison, which we would need if we use the KL divergence. We used the TD score for the sample comparisons. The TD score is easier to calculate, more comprehensible than the KL divergence, and identifies more mutations.

### Sites with significant signal change could hint to potential modifications

For the classification of mutations and modification, we compared for each site the genomic context in a 7-mer. Fig. 5 shows the absolute mean difference and average standard deviation from Eqn. 1 of each site as one data point for the `Magnipore` comparison of B.1.1.7^a^ Alpha and B.1.617.2 Delta. Most of the mismatches (blue data points) reside in the significant (green) area of `Magnipore`, as different bases result in different pA measurements. Most sites with a matching reference (orange data points) are insignificant (red background) because the same bases show similar signals and should stay consistent. However, some matching nucleotides show a significant signal difference suggesting potential differential modifications between the samples. We filtered for the coverage to ensure that inaccurate signal distributions do not cause these differential signals. A higher coverage ensures that our signal distributions have a higher data basis, so they tend to resemble the true signal distribution.

**Figure 5:**
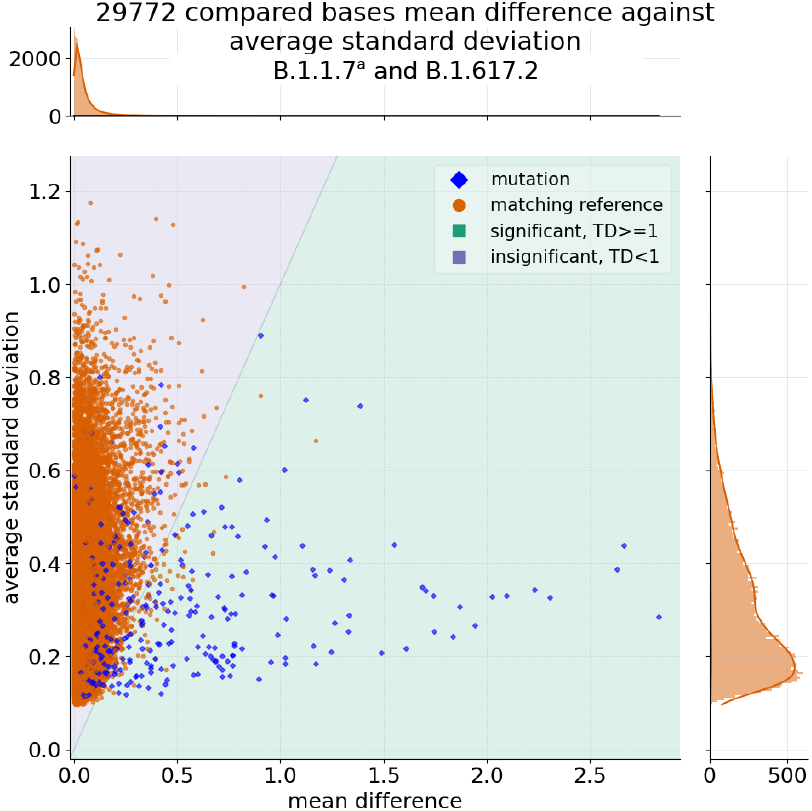
Position-wise signal comparison results between B.1.1.7^a^ Alpha and B.1.617.2 Delta. For the 29 772 compared positions the absolute mean difference is plotted against the average standard deviation. Orange – nucleotides match; blue – nucleotides mismatch. The data is not filtered for the coverage. The green and red backgrounds mark significant (signals divergent) and insignificant signal changes (signal distribution overlaps), respectively, according to Eqn. 1. The line that splits the green and red area marks the significance threshold created by Eqn. 1.

#### Low coverage leads to inaccurate signal distributions

The more coverage is given, the more data is used to approximate the signal distribution, and thus the Gaussian distribution has higher confidence to resemble the original signal distribution. Therefore, the coverage is one major factor for the comparison quality in `Magnipore`. Sites with low coverage have little data for the approximation of the signal distribution. SFig. S6 demonstrates the coverage drop in the region of ORF1a and ORF1b [11]. A low coverage can lead to skewed distributions that do not resemble the true signal distribution. When we execute `Magnipore` and look at all results without a coverage filter, we see a high amount of potential modification sites, especially in low coverage regions around ORF1a and ORF1b, SFig. S6 and SFig. S7. The number of potentially modified sites increases strongly in the Omicron samples when we do not use a coverage filter, SFig. S7. This results from inaccurate approximated signal distributions due to the low coverage.

#### Modifications can appear as mutations in reference sequences

`Magnipore` and `nanopolish eventalign` assume that the reference sequences are error-free, but this is not always true. In the ARTIC protocol RNA is converted into cDNA by reverse transcriptase. A possible modification of the RNA could introduce systematic errors in the cDNA in which only the nucleotides A, C, G, and T can be incorporated [46]. The ARTIC protocol uses the LunaScript (R) Reverse Transcriptase. According to the manufacturer, this enzyme operates at 55-65°C to melt strong secondary structures. Therefore, it is also able to transcribe areas with pronounced secondary structures. Systematic errors in the case of modified RNAs are not described. However, the introduced errors could lead to mutations between the reference sequences that do not exist in the biological sample of the ONT data [46, 47].

### Potential modification sites found repeatedly could represent variant-defining sites

We looped over all B.1.1.7^a^ comparisons and counted the potentially modified sites, Tab. 4. Interestingly, potential modification sites consistently reappear in multiple comparisons of the same sample, Fig. 6 and Tab. 4. Such sites could hint at variant or group-defining modifications, as these sites consistently differ between one sample and many others. A generally high density of potential differential modification sites can be observed in the genomic range from about 7 000 to 16 000 (ORFs: nsp 4-10 and RdRp, Fig. 6). These sites must be viewed cautiously, as they are in the low coverage range, which could lead to false positives.

**Table 4:**
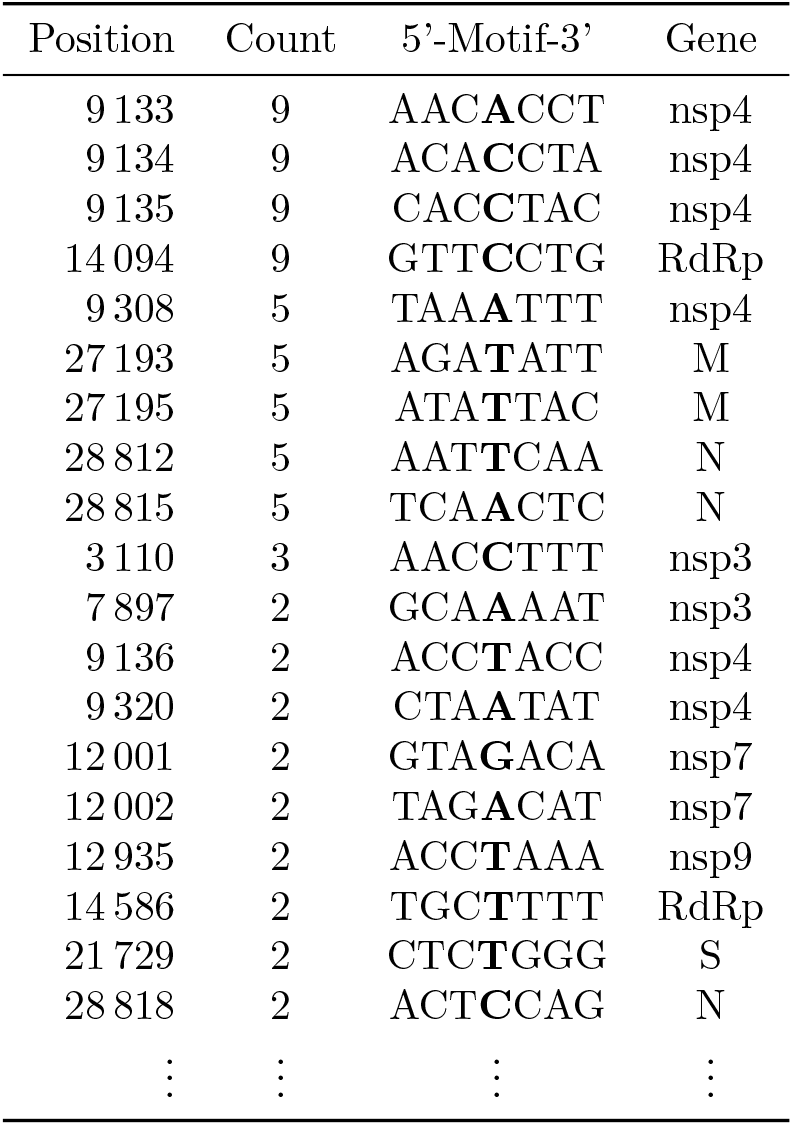
Reappearing consistent potential modification sites in the comparisons with B.1.1.7^a^ Alpha. We count them and show a part of all sites that appeared at least twice in this table. These sites must be considered in relation to the B.1.1.7^a^ reference. Therefore, we show them on the basis of their position in this strain. All underlying data can be found as CSV files in the OSF database. RdRp – RNA-dependent RNA polymerase; nsp – nonstructural protein; N – nucleocapsid phosphoprotein; M – membrane glycoprotein; S – surface glycoprotein

**Figure 6:**
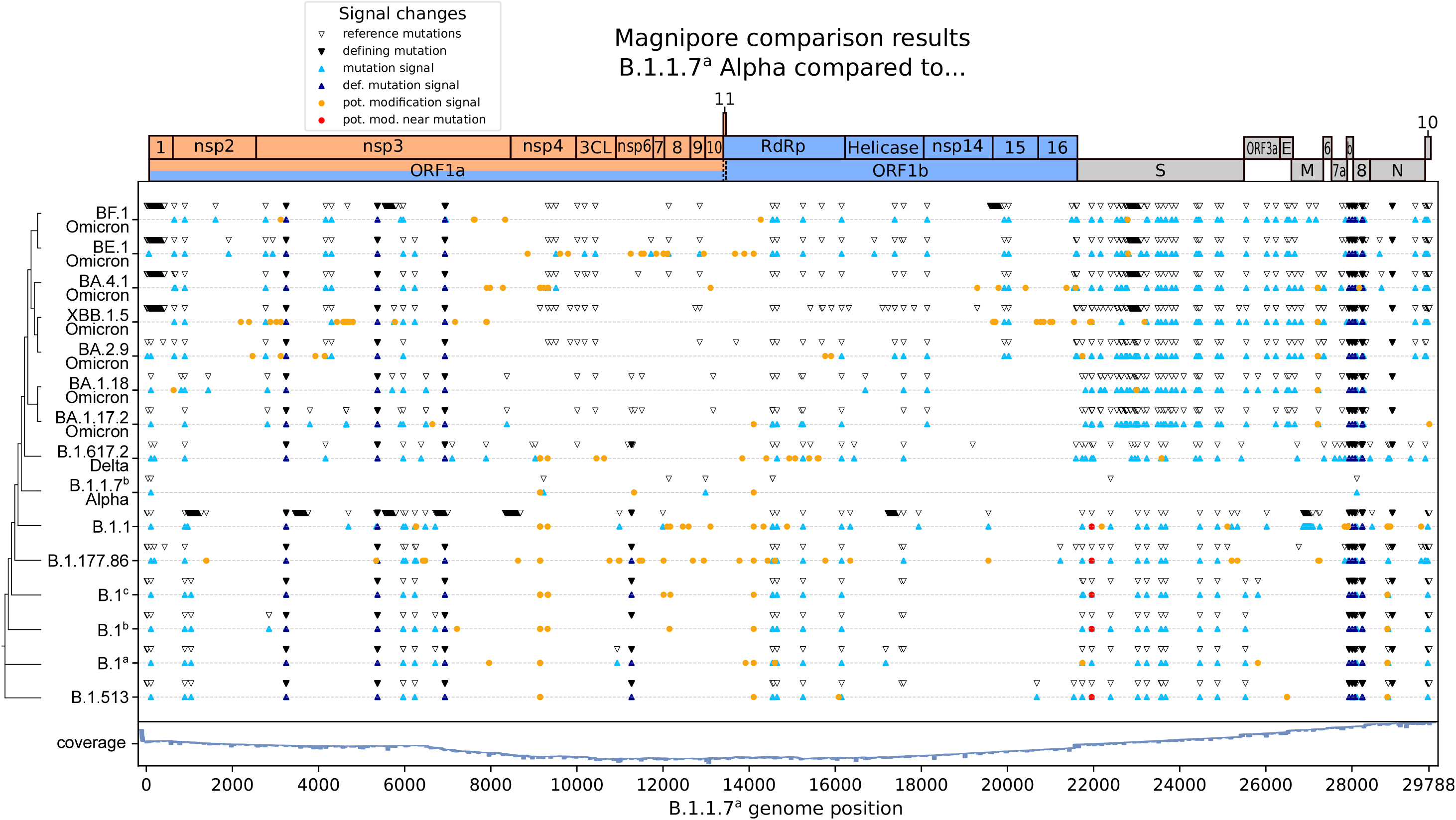
`Magnipore` sites of SARS-CoV-2 B.1.1.7^a^ Alpha comparisons to all other samples listed in Tab. 1. We marked the significant sites with a coverage of 10 or more reads on the genome of B.1.1.7^a^. For each comparison, we add two lines: the upper one containing genome information and the lower one with `Magnipore` results. White – mismatches in genome including ‘N’ regions; black – strain defining mutations for this SARS-CoV-2 strain compared to B.1.1.7^a^ taken from the GitHub repository cov-lineages (https://github.com/cov-lineages/); light blue – significant site of `Magnipore`; dark blue – significant site of `Magnipore` in agreement with defining mutations; yellow – potential modification site; red – potential modification site near repeat deletion, see Tab. 3.

#### Potential differential modification sites separate pre-Omicron and Omicron

Sites 9 133, 9 134, and 9 135 (ORF: nsp 4 Fig. 6) in the B.1.1.7^a^ Alpha genome reappear consistently across multiple comparisons. These potential differential modification sites separate pre-Omicron strains from Omicron strains. In B.1.1.7^b^ comparisons, these sites do not appear as in B.1.1.7^a^. The source for the significant signal changes is only present in the B.1.1.7^a^ comparisons.

#### Potential differential modification sites separate Omicron sublineages BA.1, BA.2 and BA.4

Contrarily, the significant sites 27 193 and 27 195 (high coverage region; ORF: M, Fig. 6) appear between B.1.1.7^a^ and of the Omicron samples: BA.1.17.2, BA.1.18, BA.2.9, BA.4.1, and XBB.1.5 (recombinant strain derived from BA.2). These sites can also be found in comparisons between B.1.1.7^b^ and the mentioned Omicron samples. They align with the same sites. When comparing both Alpha strains with the BA.5 sublineages BE.1 and BF.1, 27 193 and 27 195 do not appear as significant. Therefore, these sites could separate BA.1, BA.2, and BA.4 from the rest.

#### Further investigation is needed to clarify where the signal change is coming from

The signal distributions at 27 193 and 27 195, see SFig. S5, show the distributions of the Alpha and Omicron BA.5 samples to be very similar and tend to be bimodal (two peaks). In contrast, the other Omicron distributions are visibly different and unimodal (single peak). The source of the differential signal between these groups of variants must be investigated in further studies. Possible sources for a bimodal signal are (1) a small percentage of reads have a modification, (2) a small percentage of reads have a mutation, or (3) the segmentation tools struggle to segment the signal at this position accurately. 27 193, and 27 195 could indicate that the viral epitranscriptome differs at these sites characteristically. Sites 27 193 and 27 195 occur with a mutation in the samples’ references at position 27 210 in the B.1.1.7^a^ genome. This strain shows an ‘A’, whereas the Omicron variants have a ‘C’ in the alignments. Our BA.5 references do not have this mutation and do not show a potential differential modification. The ‘A/C’ mutation is 24 nucleotides downstream from the potential differential modification. Hence, the nucleotides are not measured together in the pore, and the mutation should not directly influence the measured electric current.

Furthermore, as the mutation is downstream in the RNA sequence, it will be sequenced before the position of the potential differential modification. The mutation has left the pore by the time the potential differential modification is sequenced. Intrinsic properties of the RNA, such as secondary or tertiary structures and RNA modifications, influence the translocation process and time (helicase stalling) [48, 49], but not the intensity of the electric current.

#### Potential differential modification sites separate variants of concern

Sites 28 812 and 28 815 (ORF: N, Fig. 6) seem to separate the variants of concern (VOCs: Alpha, Delta, Omicron) from the rest (except B.1.177.86). Comparisons with more pre-VOC samples are necessary to substantiate this finding. We only have samples for 4 different pre-VOCs; three B.1 samples could bias this result.

#### All potential differential modification sites can be found in the OSF database

To identify more defining modification candidates, we looped over all comparisons and collected potentially modified sites that appeared in at least three comparisons, Fig. 7. These sites are listed in the OSF database.

**Figure 7:**
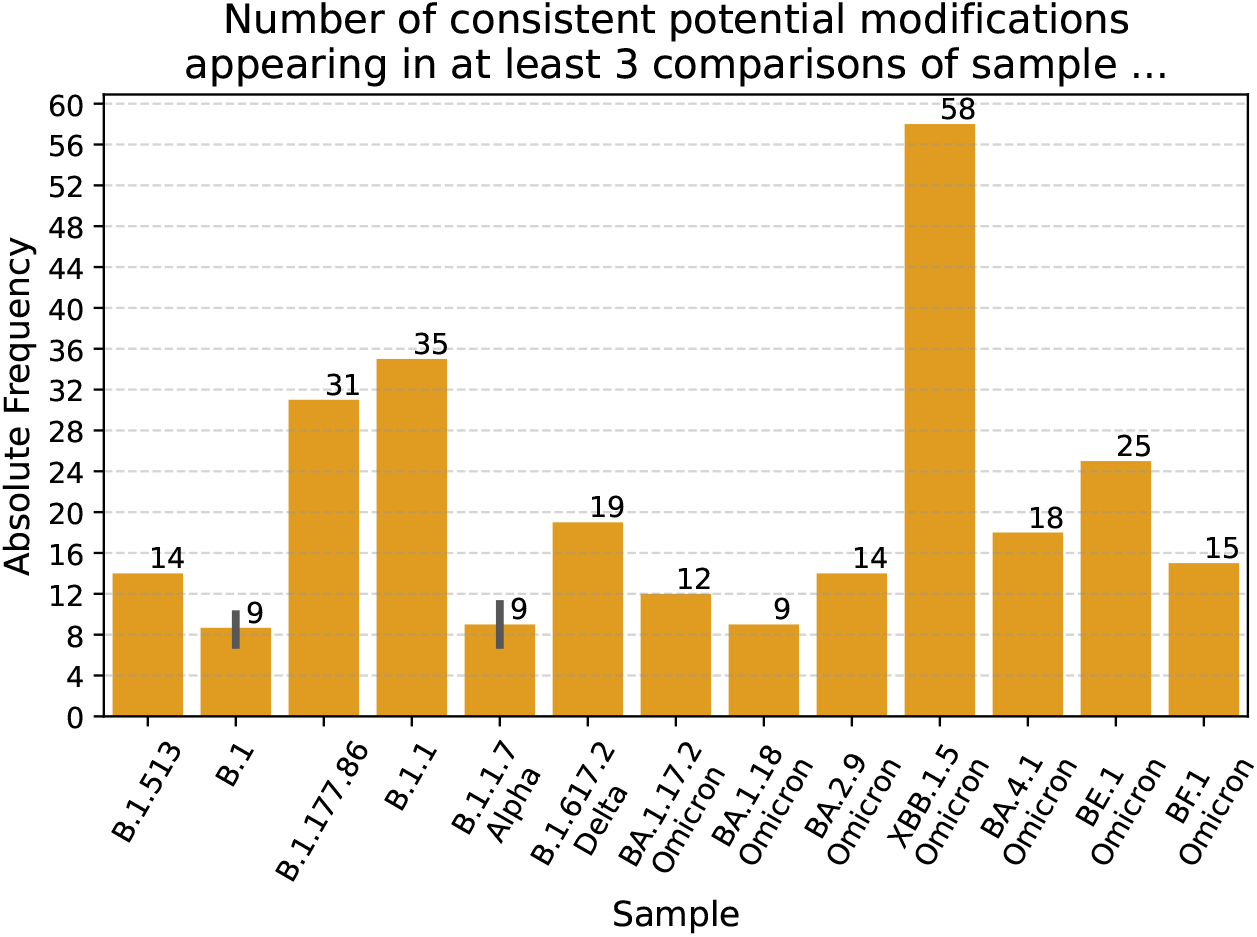
Consistently reappearing RNA modifications. For each site in each sample comparison against all other samples of Tab. 1, the number of significant sites classified by `Magnipore` as potentially modified is given. If a site appears in three or more sample comparisons as potentially modified, then we count this site as a consistently reappearing modification. These sites should be investigated further with higher priority as they might hint at potential defining modifications. A list of all sites can be found in the OSF database.

### Knowledge of the localization of RNA modifications may be useful to optimize vaccines or search for antiviral targets

COVID-19 mRNA vaccines use, among others, the modified nucleobase N1-methylpseudouridine to reduce RNA immunogenicity and increase translational efficiency [50]. Knowledge of the location and nature of RNA modifications may also be important for the development of therapeutic approaches, especially since such modifications have been described in a wide range of RNA viruses [14, 15, 16, 17, 18].

#### False positive potential modifications – limitation by nanopolish eventalign

One source for deviating signals at potential modification sites could be mutated subsets of viruses called haplotypes. Mutations get erroneously corrected as `nanopolish eventalign` tries to correct all reads according to the provided reference sequence. Signals caused by mutations in individual reads might be assigned to the wrong bases. The mutation signals then shift and skew the signal distributions at these sites, SFig. S4, which could lead to false positive potential modification sites between samples. Mistaking mutations for basecalling errors is a limitation of the resquiggle tool.

The exact signal distributions for each potential modification must be investigated individually to further investigate possible false positive sites, SFig. S9.

#### Possible influence of virus cultivation and RNA preparation techniques on the prediction of modification sites

Another reason for the observed variability could be the virus cultivation and the subsequent RNA preparations. For the propagation of SARS-CoV-2, commercially available Vero cells were used, originating from the kidney of African green monkeys [51]. It is known from experiments with arboviruses that the cell line itself influences RNA modifications. Thus, the modifications differ between Sindbis virions that were propagated in insect cells and those from mammalian cells. Virions propagated in the insect cells replicated better in the mammalian cells [52]. Although certainly not as pronounced as in the example mentioned above of a switch between insect and mammalian cells, it is conceivable that replication of SARS-CoV-2 in other mammalian cells could impact the modification pattern and, thus, the prediction of modified sites in the genome. It is also possible that the modifications of the viral RNA from the cell are different from those of the infectious virion. The RNAs we sequenced originate mainly from cell lysates but may also contain portions of extracellular virions. Another aspect is the occurrence of subpopulations *in vivo* and *in vitro*, so-called quasi-species [53], which, as explained in the previous section, can influence the results. In addition, the RNA was prepared after the occurrence of a cytopathic effect detectable by light microscopy. Standardization to the virus dose used and the cell cycle did not take place. This could be taken into account in future experiments. Likewise, different host cells and RNA preparations from purified virions of the culture supernatant should be investigated.

### Modifications tend to occur in stacks of predicted RNA secondary structures

Previous studies have shown that the RNA of SARS-CoV-2, like that of other betacoronaviruses, has conserved secondary and probably tertiary structures essential for the viral life cycle [54, 55, 56, 57, 11]. We investigated the correlation between modifications and *in silico* predicted RNA secondary structures in genomes of betacoronaviruses (including SARS-CoV-2).

`Magnipore` identified eight differentially modified sites across the entire genome which occur in at least 40% of all pairwise comparisons: 2 473, 3 932, 4 148, 15 915, 15 920, 16 143, and 27 211, 27 213 (position in BA.2.9 genome). We detected six potential differentially modified sites in a stack of base pairs, Fig. 8. The remaining two sites (15 920 and 16 143) are located in a hairpin loop and an internal loop, respectively. The alignments reveal a high sequence similarity resulting in predicted base pairs only containing one base pair type (labeled red in alignment and secondary structure in Fig. 8). Three modification sites are located in ORF1a. We predicted structural elements in the nonstructural protein (nsp) 2 gene containing a small hairpin loop with only three base pairs and a 47 nt long secondary structure with two internal loops (see Fig. 8, left bottom). This predicted formation molecule has a p-value of 2.5 *×* 10^*−*3^. One modification in the nsp3 gene is located in an RNA secondary structure consisting of three stem-loops (one with an internal loop). We observed the lowest p-value (6.94 *×* 10^*−*23^) compared to the other sequence-structure alignments in this RNA molecule in this study. According to our computational predictions, the other modification site in the nsp3 gene may be located in a multi-loop. We observed more divergent base pairs within this structure according to the possible base pair types. Even two base pairs occur with three or four types (green and cyan colored in Fig. 8 top middle). We predicted two structures in ORF1b in the RNA-dependent RNA polymerase (RdRp) gene. The first element includes two modification sites, one in a stack and the other in the hairpin. This prediction shows the highest p-value (1.86 *×* 10^*−*2^) and many predicted base pairs do not fold in two or more sequences of the alignment, assuming that this structure might not be present. The second element in the RdRp gene revealed a molecule of three stem-loops (one with several internal loops). The high level of sequence conservation in this region leads to a high number of base pairs with only one base pair type (red). However, within this RNA secondary structure, we observed three different types of basepairs (green) at one position, giving a higher confidence in the existence of the structure. The last two modification sites are located in ORF6 and present in a stacking of base pairs (see Fig. 8 bottom right). One modification is in a base pair with one possible base pair type. The other basepair does not fold in more than two sequences in the alignment. The prediction of this region revealed three stem-loops in total with a p-value of 1.36 *×* 10^*−*17^. A pairing in the last element is possible with three base pair types.

**Figure 8:**
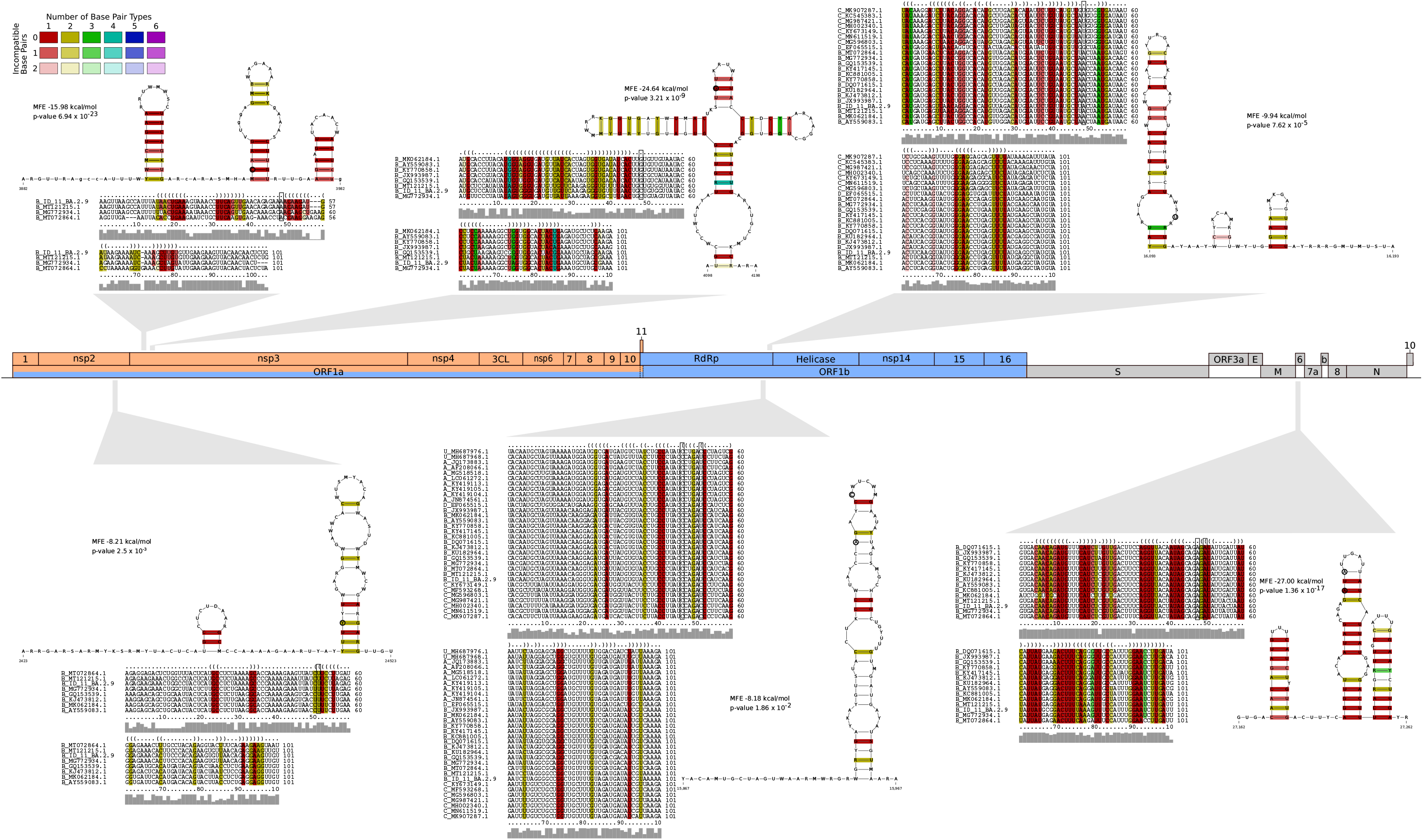
Predicted RNA secondary structures of betacoronaviruses in correlation with differentially modified sites in SARS-CoV-2 genome. Predicting structural elements by a computational alignment-based approach, we observed six out of eight sites in a stacking of base pairs. The remaining two are in a hairpin and internal loop. The sequence similarity in the respective regions is high. Thus, many predicted base pairs are only folded by one base pair type (colored red in alignment and secondary structure). Nevertheless, some predicted structural elements show three or four possible base pair types (green and cyan). The betacoronavirus lineages (A-D), according to ICTV [58], were added to the respective accession IDs in the alignments. Unclassified genomes are labeled with U.

The structures shown here were predicted using computational methods only and may not reflect secondary structures found in betacoronaviruses *in vivo*. Across the potential modification sites and predicted RNA secondary structures, we observed the tendency of differentially modified sites occurring in a stack of base pairs.

### Comparison with similar tools

We used B.1.1.7^a^ Alpha and B.1.617.2 Delta to compare `Magnipore` with `xPore`, `Nanocompore`, and `Yanocomp`. These tools work similarly to `Magnipore` by searching for differential signals within the ONT data. We executed `Magnipore` with default parameters. We used `Guppy` 6.1.7 with the `rna_r9.4.1_70bps_hac` model to basecall the SARS-CoV-2 RNA sequencing data. We use the resulting reads from `Guppy` for every tool in the comparison to establish comparable conditions. The B.1.1.7^a^ sample has 798 140 basecalled reads with around 1 308 000 000 bases, of which 1 023 000 000 could be mapped to the B.1.1.7 reference sequence. B.1.617.2 had 526 766 basecalled reads with around 67 200 000 bases, of which only 14 860 000 could be mapped to the B.1.617.2 reference sequence. We consider only sites with a coverage of at least 10 in both samples.

For sample B.1.1.7^a^ these are 29 784 of 29 788 nucleotides, and for sample B.1.617.2 25 413 of 25 413 of 29 858 bases.

#### Magnipore detects 161 significant sites consisting of 146 mutation and 15 potential modification sites

The 161 identified sites have a significant TD score greater or equal to 1 (see materials and methods). `Magnipore` classified these sites into 146 mutation and 15 potential modification sites.

Multiple adjacent sites can appear with a significant differential ONT signal caused by only one molecular modification or mutation, Fig. 2A. Likewise, the number of differing ONT signals is larger or equal to the number of naturally appearing mutations and detected mutations reported by `Magnipore`.

`Magnipore`s 146 mutation sites resulted in 110 detected mutations, of which 55 are non N mutations, and a high fraction of 55 mutations include an N at the site of interest. With this result, `Magnipore` missed to identify 20 mutations: The ground truth (read from the alignment between B.1.1.7^a^ and B.1.617.2 of the reference sequences) is 130 mutations, of which 64 are non N, and 66 are N mutations.

The 15 potential modification sites are not close to a mutation, i.e., not in the range of three nucleotides up or downstream of a mutation. At these sites, the signal significantly differs between the samples without a mutation in the reference sequence alignment. Potentially modified sites need further investigation because of software limitations like inaccurate signal segmentation or incorrect error correction by `nanopolish eventalign`.

For this comparison, `Magnipore` needed around 35 hours to process, compare, write, and plot the data of both samples on an AMD Ryzen 9 3900X 12-Core Processor with 24 threads and an NVIDIA GeForce RTX 2080 Ti. Basecalling took 1 hour and 17 minutes (B.1.1.7) and 26 minutes (B.1.617.2). Mapping and `nanopolish` preparation took 4 hours and 20 minutes (B.1.1.7) and 21 minutes (B.1.617.2), respectively. Creation of the distribution models for every position took 24 hours and 20 minutes (B.1.1.7) and 3 hours and 43 minutes (B.1.617.2). Comparing the position-wise signal distributions with the help of an alignment took just two minutes.

#### xPore misses sites and classifies mutations as modifications

`xPore` analyzes only 2 057 sites and mistakes mutations for modifications. As preparation for `xPore`, both samples are mapped to only one reference and run `nanopolish eventalign`. We map both samples to the B.1.1.7^a^ Alpha reference. For `nanopolish`, the parameters signal-index and scale-events are used. We execute `xpore` dataprep with readcount_min 10 to filter for sites with a coverage of at least 10, compare the output to `Magnipore` and prepare the data for further analysis with `xpore`. We execute `xpore` diffmod with default parameters and compare the output to `Magnipore`. `xPore` provides results for 2 057 positions in the genomic range (not every position present in the ranges) of 2 to 1 710 nt and 25 347 to 29 783 nt on the B.1.1.7^a^ genome, which is roughly 6.9% of all positions. For further analysis, we filtered these 2 057 sites for a p-value below 0.01 and reduced the set to 553.

`Magnipore` and `xPore` share 23 significant sites with a p-value below 0.01 by `xPore` and a TD score higher or equal to 1 by `Magnipore` (sixth bar in Fig. 9). `xPore` does not differentiate between modification and mutation sites, as it expects to compare the same biological sample in two different conditions. It only reports differential modification rates. `xPore` reports 48 significant sites out of the 553 in close proximity to 25 mutations (fifth bar in Fig. 9). Therefore, `xPore` mistakes 48 mutations as significantly differentially modified. These are 48 false positive sites regarding `xPore`s task to find differential modification. `Magnipore` can compare different SARS-CoV-2 variants that contain mutations and reports them. `Magnipore` detects 161 significant positions. It detects and correctly classifies 55 mutations. `Magnipore` shares 23 significant sites with `xPore`, which are in close proximity to 20 mutations. `Magnipore` and `xPore` do not share potential modification sites.

**Figure 9:**
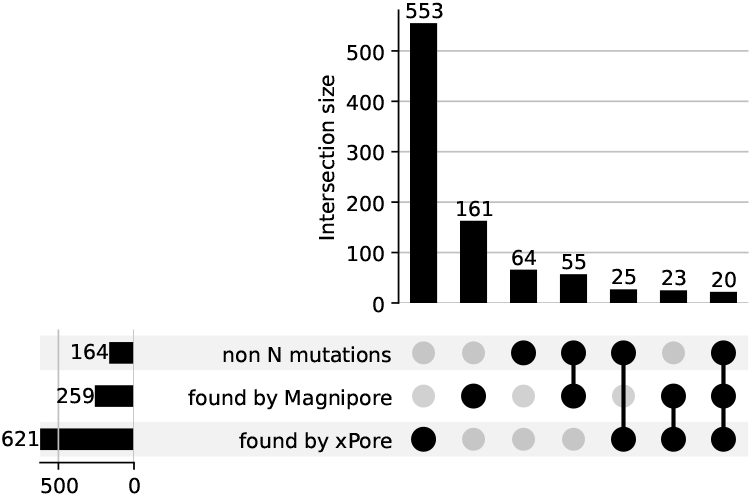
Upset plot showing sets and intersections between present non N mutations, found positions by `Magnipore` and found positions by `xPore`. We compared the output of `Magnipore`, `xPore`, and the set of present mutations between B.1.1.7^a^ Alpha and B.1.617.2 Delta. Connected dots indicate set intersections. Bars show the number of elements (positions) in this intersection. Intersections with non N mutations represent the number of identified mutations by the respective tool.

For `Nanocompore`, we followed the guidelines in their documentation. As `Nanocompore` only works on a single reference genome, we mapped the reads from the B.1.617.2 sample to the B.1.1.7^a^ reference sequence. We used the resulting mappings to execute `nanopolish eventalign` for both samples with the required parameters `--print-read-names --scale-events --samples`. After that, we ran `nanocompore` eventalign_collapse on both samples with default parameters and executed `nanocompore` sampcomp on the output files. We could not run `nanocompore` sampcomp without getting an error.

We ran `yanocomp` according to the guideline on https://github.com/bartongroup/yanocomp. The data was prepared with `nanopolish eventalign` using the parameters `--print-read-names --scale-events and --signal-index`. We executed `yanocomp` prep with default parameters. Finally, we used `yanocomp gmmtest` with default parameters and `--min-read-depth 10`. We could not get any result with `yanocomp` and stopped `gmmtest` after 10 days of running.

## Conclusion

### Limitations

Further improvement to `Magnipore` and its signal analysis capabilities will require a better resquiggling and segmentation algorithm of `nanopolish eventalign`. An essential task in the future will be a precise differentiation between basecalling errors, mutations, and modifications among the reads and the reference sequence of the sample. Another way to confirm the results of `Magnipore` would be to have multiple samples or replicates from the same variant. This way, we could gain confidence about consistent potential modification signals.

Multiple aspects influence the analysis of two related samples with `Magnipore`. One is the quality of the reference sequence. A high-quality reference sequence (1) should not contain ambiguous bases (N), (2) should not miss bases at the 5’ and 3’ endings, and (3) should not contain wrong bases (errors). `Magnipore` can also work with related reference sequences, but this will reduce the quality of the results. `Magnipore` depends heavily on the segmentation and resquiggling quality of `nanopolish eventalign`. Improvements in resquiggling and segmentation of the ONT signal would be a significant step forward in the performance of `Magnipore`, as the range of influence from mutations can be narrowed down. An evenly distributed high coverage or deep sequencing is optimal for the best analysis. `Magnipore` analyzes the raw ONT signal and only depends on the `Guppy` basecaller to map the reads to the corresponding reference sequence for the segmentation step. Other steps are independent of the basecaller. The tool can find sites with a significant signal change on a genomic level that appear consistently in different sample comparisons.

### Outlook

With the help of RNA sequencing and `Magnipore`, nucleotide positions that may be differentially modified can be identified. Individual sites are found across variants, while some appear specific to groups of more closely related variants. However, these are only predictions so far, and there is no experimental confirmation yet of the suspected differential RNA modifications between SARS-CoV-2 variants. From our point of view, `Magnipore` detects promising sites for potential differential RNA modifications and, in some cases, even potential defining RNA modification sites.

Findings about the localization and type of chemical modifications in the SARS-CoV-2 genome could serve as a starting point for advancing antiviral drug discovery. It is also conceivable that this knowledge will help improve the effectiveness of existing mRNA vaccines.

## Acknowledgements

The authors gratefully acknowledge the help of Sebastian Krautwurst in understanding and handling Oxford Nanopore data. The authors thank Dr. Barbara Mühlemann (Institute of Virology, Charité Berlin) for providing the Illumina reference sequences. Special thanks go to Dr. Thomas Lorentz (Labor Dr. Krause) and Prof. Helmut Fickenscher (Institute for Infection Medicine) for providing the equipment, reagents and facilities.

## Author contributions

Provided the SARS-CoV-2 RNAs and ARTIC-protocol generated reference sequences: AK, MMH, CMC, LP, AW, RR; RNA sequencing: MZ, AS; Tool development: JS; Algorithm design: JS, CHzS, MM; Wrote the paper: JS, CHzS, MM, MZ, AK, ST; Guided the study: CHzS, AK, MM.

## Code availability

The `Magnipore` pipeline is written in *python3* and available via the GitHub repository https://github.com/JannesSP/magnipore and via Conda https://anaconda.org/JannesSP/magnipore.

## Data availability

The raw ONT sequencing data and basecalled `FASTQ` files can be found in the SRA database under the BioProject: [PR-JNA907180](https://dataview.ncbi.nlm.nih.gov/object/PRJNA907180?reviewer=pgvdc7697efvrt1p19qek2em3d). The output of `Magnipore`, the list of consistently reappearing potential modified positions and reference sequences can be found in the OSF database https://osf.io/evc6k/.

## Footnotes

1 https://github.com/klamkiew/cov_trs_structure/blob/master/diNuclShuffle.py

