## Supplemental Material for "Magnipore: Prediction of differential single nucleotide changes in the Oxford Nanopore Technologies sequencing signal of SARS-CoV-2 samples"

### Supplemental Materials

Table S1: The presented samples are stored together with an identifier within the SRA und OSF databases. The corresponding IDs can be found in this table. B.1.1.7[a], B.1.1.7[b], B.1.617.2, BA.1.17.2, BA.1.18, and BA.2.9 were additionally sequenced by the Institute of Virology in the Charité Berlin on the Illumina platform MiSeq using the ARTIC protocol to assemble reference sequences using `bowtie2` [59]. The Illumina-assembled references except BA.1.18 can be found in the GISAID database [60] using the provided IDs from Tab. 1. The remaining references were sequenced and assembled as described in Sec. Preparation of total RNA. All reference sequences are stored in the OSF database (`https://osf.io/evc6k/`) together with additional material and plots. The raw ONT sequencing data and basecalls are stored in the SRA database (BioProject: PRJNA907180).

| Samples | ID in databases |
|---|---|
| B.1.1.7[a] | ID 1 B.1.1.7 |
| B.1.617.2 | ID 2 B.1.617.2 |
| B.1.177.86 | ID 3 B.1.177.86 |
| B.1.1 | ID 4 B.1.1 |
| B.1[a] | ID 5 B.1 |
| B.1[b] | ID 6 B.1 |
| B.1[c] | ID 7 B.1 |
| B.1.513 | ID 8 B.1.513 |
| BA.1.17.2 | ID 9 BA.1.17.2 |
| BA.1.18 | ID 10 BA.1.18 |
| BA.2.9 | ID 11 BA.2.9 |
| BE.1 | ID 12 BE.1 |
| BF.1 | ID 13 BF.1 |
| B.1.1.7[b] | ID 14 B.1.1.7 |
| BA.4.1 | ID 16 BA.4.1 |
| XBB.1.5 | ID 18 XBB.1.5 |

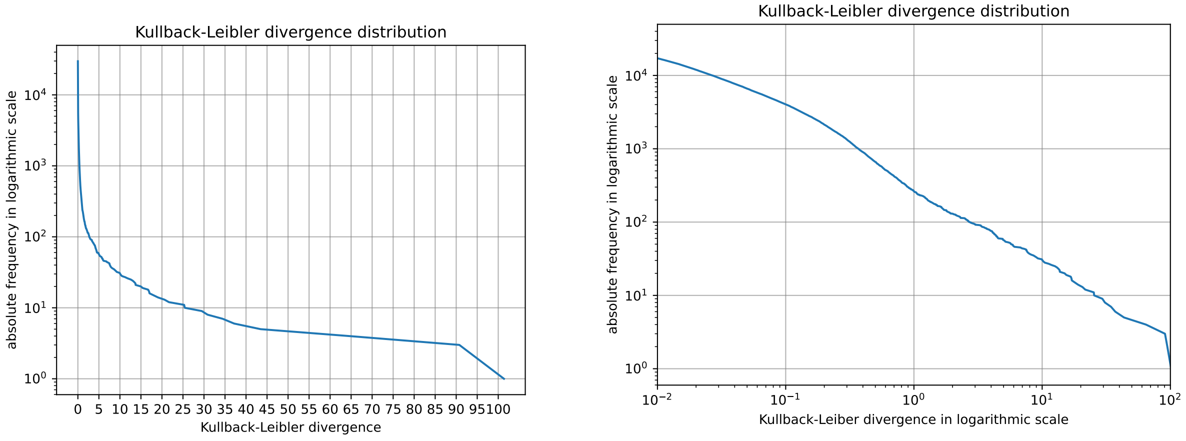


Figure S1: The distribution of the Kullback-Leibler (KL) divergence in the sample comparison between B.1.1.7[a] Alpha and B.1.617.2 Delta. Both diagrams show the KL divergence value for each compared site on the x-axis in default scale **left** and logarithmic scale **right**. The y-axis shows the absolute frequency in logarithmic scale in both diagrams. The comparison of TD score results with the results when we use a KL divergence threshold are given. By analyzing the KL divergence distribution we decided for a threshold 1, as we see 29510 sites with a KL divergence value below 1. In the **left** figure we see, that the KL divergence value higher 1 is distributed more uniformly.

**Illumina references have less ambiguous bases than ONT references.** For samples B.1.1.7[b] Alpha, B.1.617.2 Delta, BA.1.17.2 Omicron, BA.1.18 Omicron, and BA.2.9 Omicron we compared the reference sequences assembled from Illumina data by the Institute of Virology in the Charité Berlin to the reference sequences assembled from ONT MinION data. The Illumina references have more unambiguous bases than the ONT references which is why we used the Illumina references for the `Magnipore` analysis if we had one.

Table S2: In all cases the reference sequence assembled from the ONT data is longer than the references assembled from the Illumina data. But the ONT references have much more N content and therefore less unambiguous characters than the Illumina references.

| Sample | Illumina | #N | AT : GC : N | ONT | #N | AT : GC : N |
|---|---|---|---|---|---|---|
| B.1.1.7[b] | 29760 | 2 | 0.620 : 0.380 : 0.000 | 29885 | 201 | 0.616 : 0.377 : 0.007 |
| B.1.617.2 | 29858 | 0 | 0.620 : 0.380 : 0.000 | 29890 | 199 | 0.616 : 0.377 : 0.007 |
| BA.1.17.2 | 29756 | 64 | 0.619 : 0.379 : 0.002 | 29874 | 751 | 0.605 : 0.370 : 0.025 |
| BA.1.18 | 29830 | 3 | 0.620 : 0.380 : 0.000 | 29873 | 157 | 0.617 : 0.378 : 0.005 |
| BA.2.9 | 29808 | 2 | 0.621 : 0.379 : 0.000 | 29850 | 720 | 0.605 : 0.371 : 0.024 |

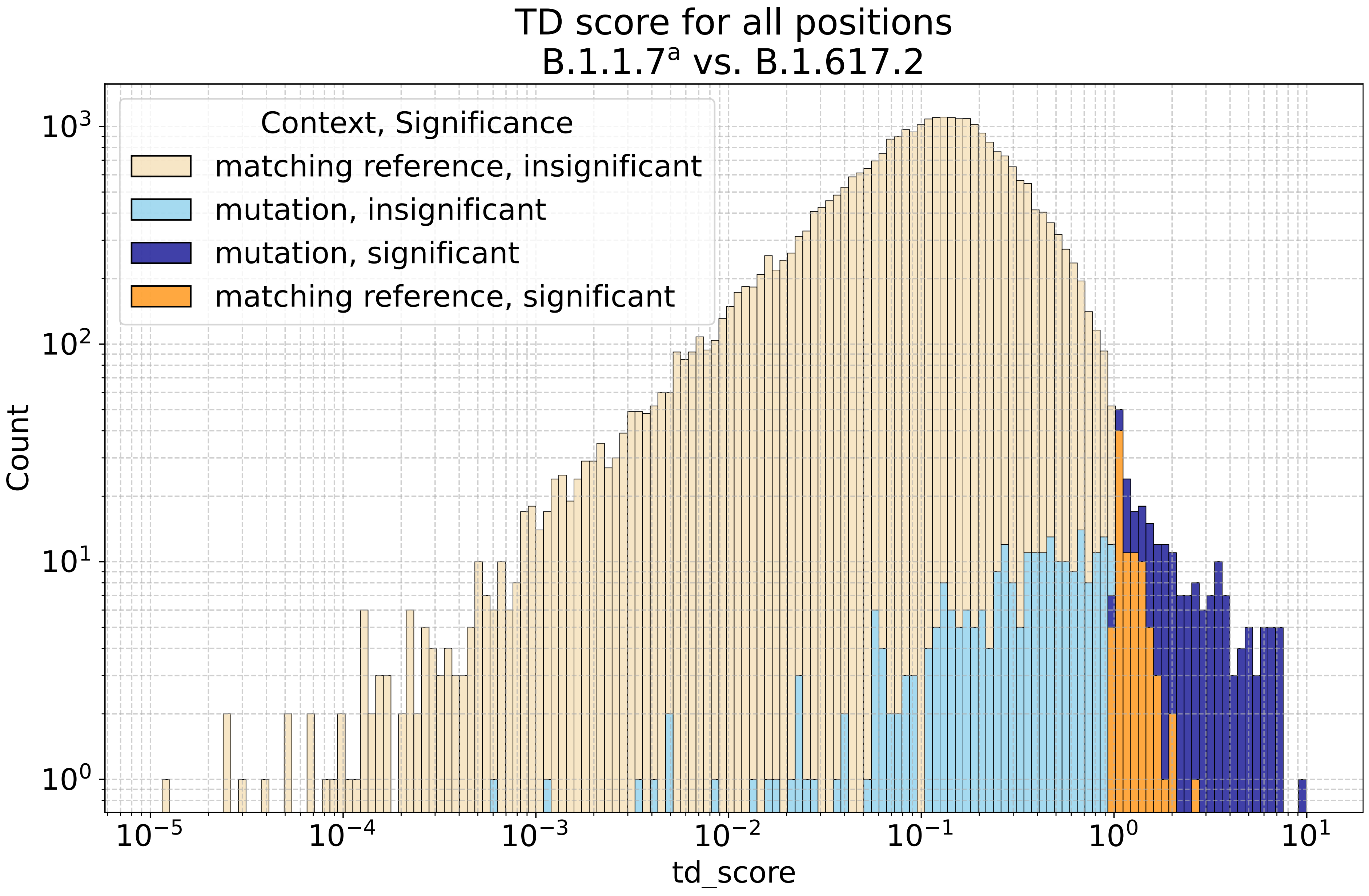


Figure S2: Threshold distance (TD) score distribution across the sites. Mutations tend to have a higher TD score than matching sites between B.1.1.7[a] Alpha and B.1.617.2 Delta. On the x-axis we can observe the TD score in logarithmic scale. A TD score of 1 ($10^0$) marks the significance threshold. The y-axis shows the counts of sites with the corresponding TD score also in logarithmic scale. Light orange – matching bases in the reference with an insignificant TD score; light blue – sites that show a mutation between the samples and have an insignificant TD score; dark blue – mutation sites with a significant TD score; dark orange – matching bases with a significant TD score.

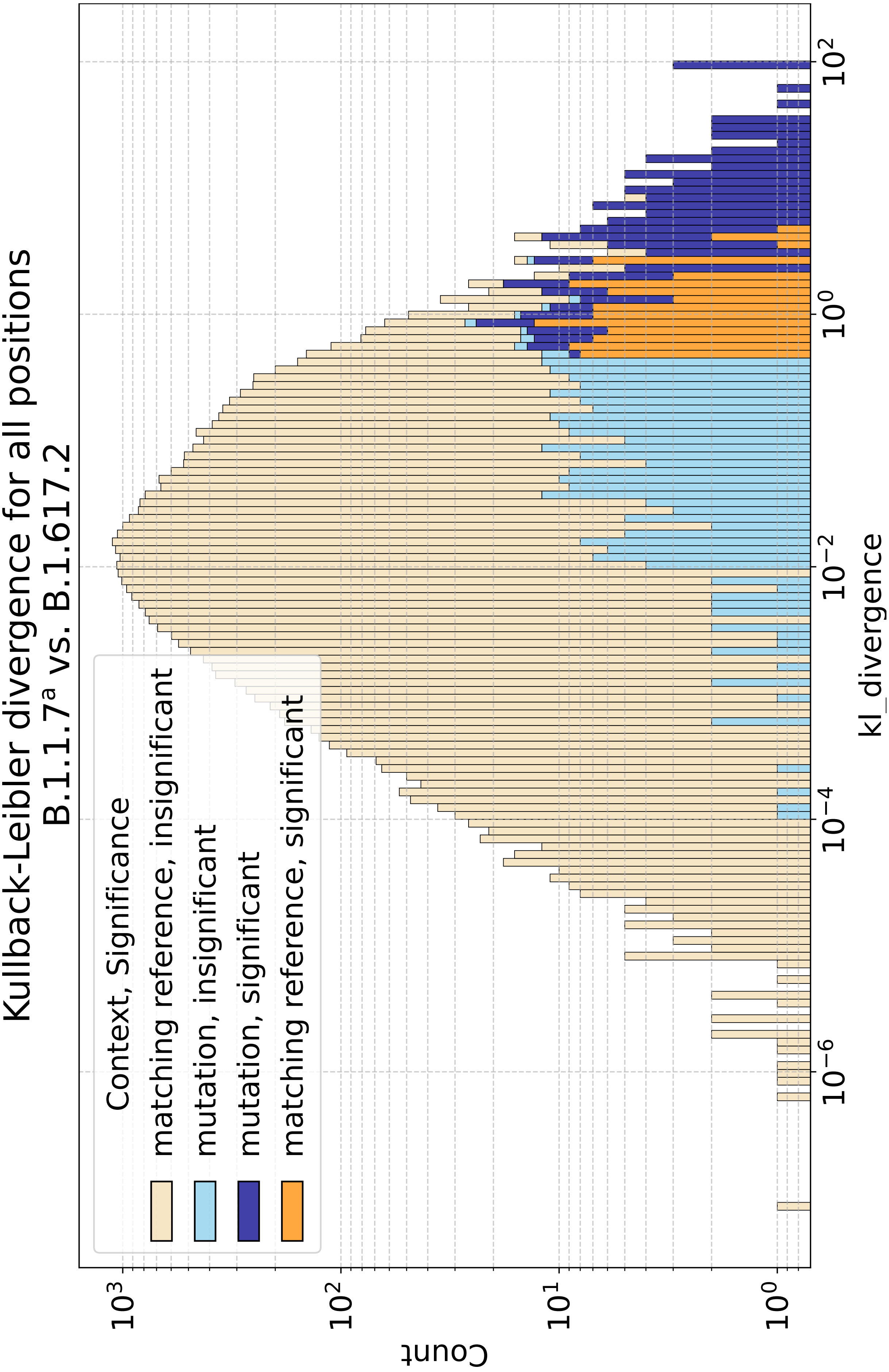


Figure S3: The similarity of the Kullback-Leibler (KL) divergence to the threshold distance (TD) score in Fig. S2. The data is also colored the same as in Fig. S2. The KL divergence shows a similar distribution as the TD score. If we choose a threshold of 1 KL divergence we detect less mutations compared to the TD score. We also have less significant sites with a matching reference that might hint to differential modifications.

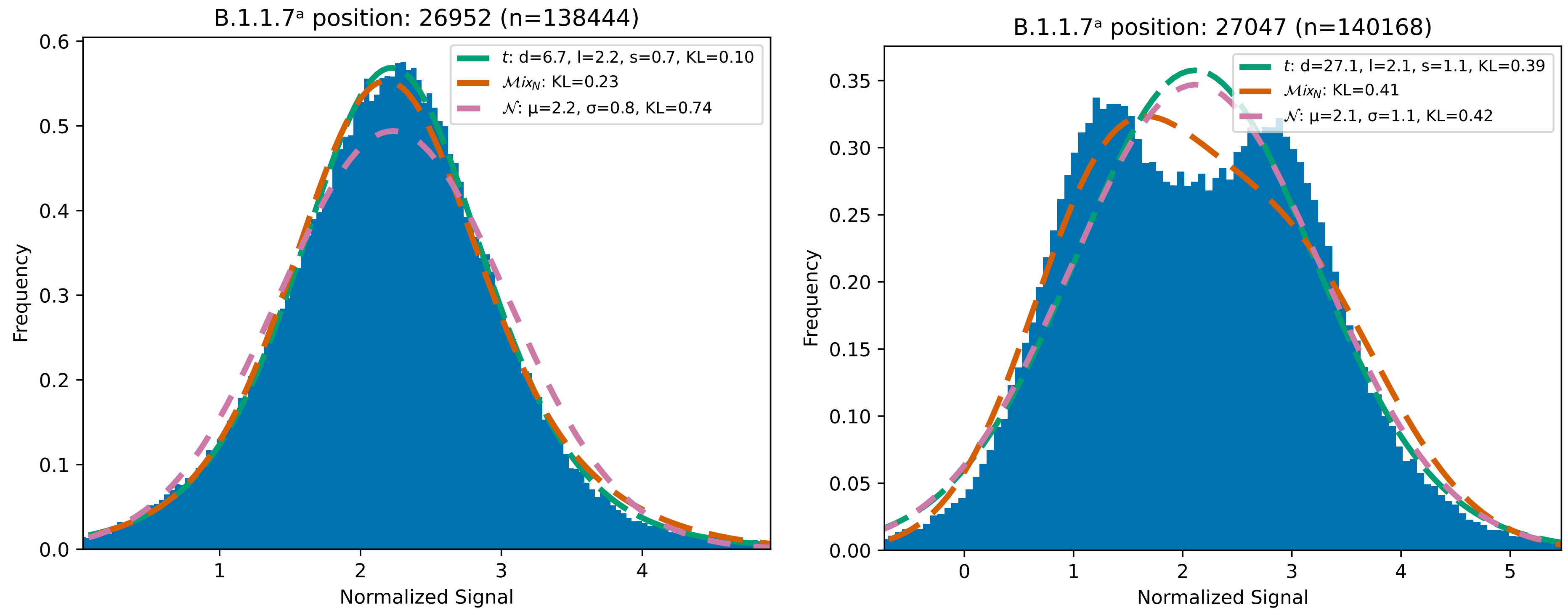


Figure S4: The Gaussian distribution is a good approximation for a signal distribution per position in case of ONT sequencing. But there are other more complex distributions that fit better, as seen in the KL divergence value in the legend (lower is a better fit). We basecalled the raw ONT data and mapped the reads against their corresponding reference (B.1.1.7[a] Alpha or B.1.617.2 Delta). We then resquiggled and segmented the data with `nanopolish eventalign`. For all segments (n), the read normalized mean is plotted in the histogram. **Left**: For many signal distributions the Gaussian distribution is a good approximation. More complex distribution like the Students-t or Gaussian mixture fit much better. **Right**: Some positions show a bimodal signal distribution. A reason for that could be inaccurate segmentation by `nanopolish eventalign` or some reads are modified or carry a mutation.

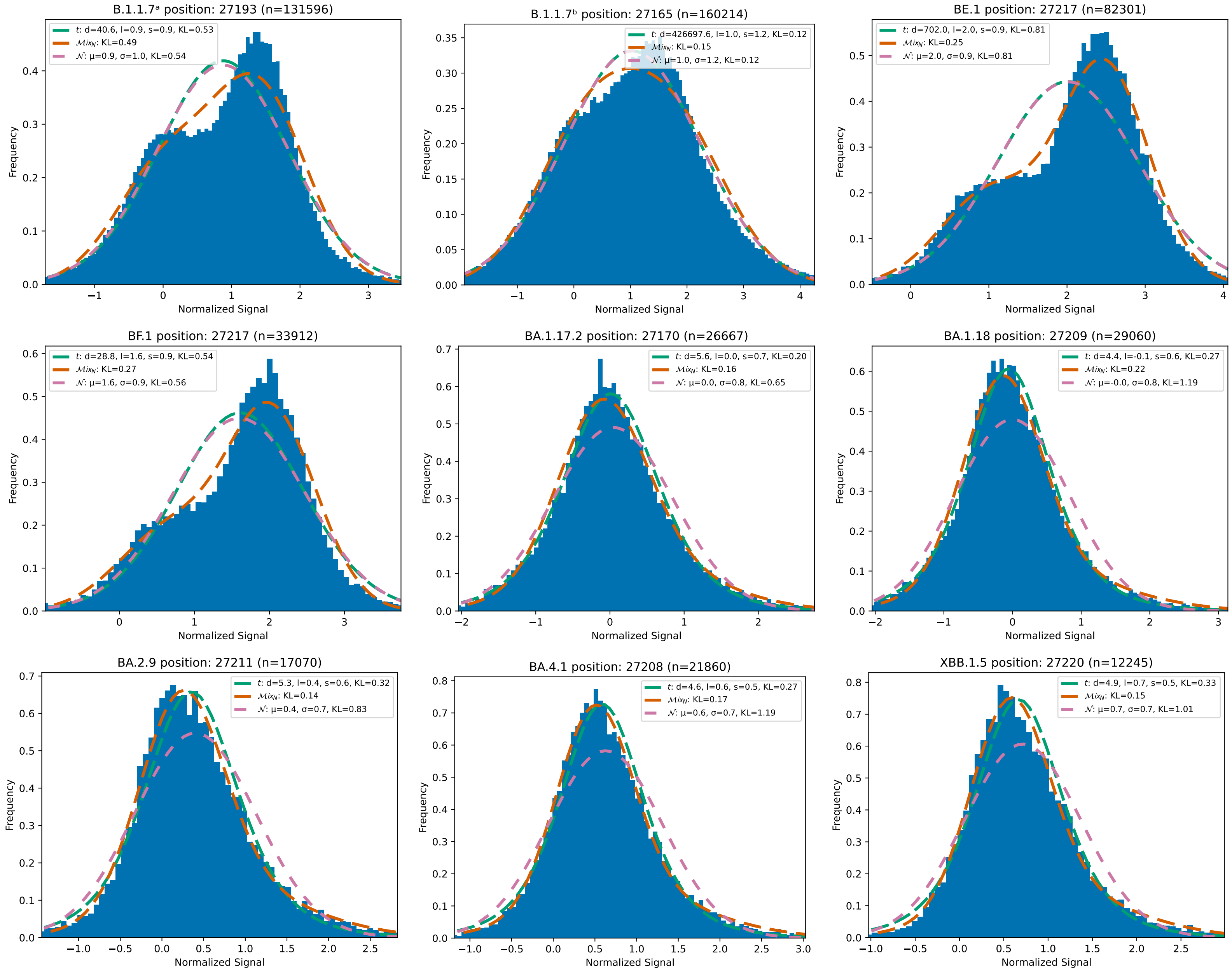


Figure S5: Segment mean distributions for all Alpha and Omicron samples: `Magnipore` detects potential differentially modified sites between the set of B.1.1.7[a], B.1.1.7[b], BE.1, BF.1 and the set of BA.1.17.2, BA.1.18, BA.2.9, BA.4.1, XBB.1.5. These sites are promising defining modification sites, separating BA.1, BA.2, and BA.4 from all other variants in our study. The signal distributions of the Alpha and BA.5 samples tend to be bimodal, while the others are unimodal. The figures Further investigations are needed to find the source of the bimodality. Possible reasons could be (1) modified reads, (2) mutated reads, or (3) inaccurate signal segmentation. Three distributions are fitted: $\mathcal{N}$: a Normal distribution with mean $\mu$ and standard deviation $\sigma$; $Mix_{\mathcal{N}}$: a mixture of two Normal distributions; $t$: a Students'-t distribution with $d$ degrees of freedom, location $l$, and scale $s$. For each distribution, the Kullback-Leibler (KL) divergence from the histogram is given as well.

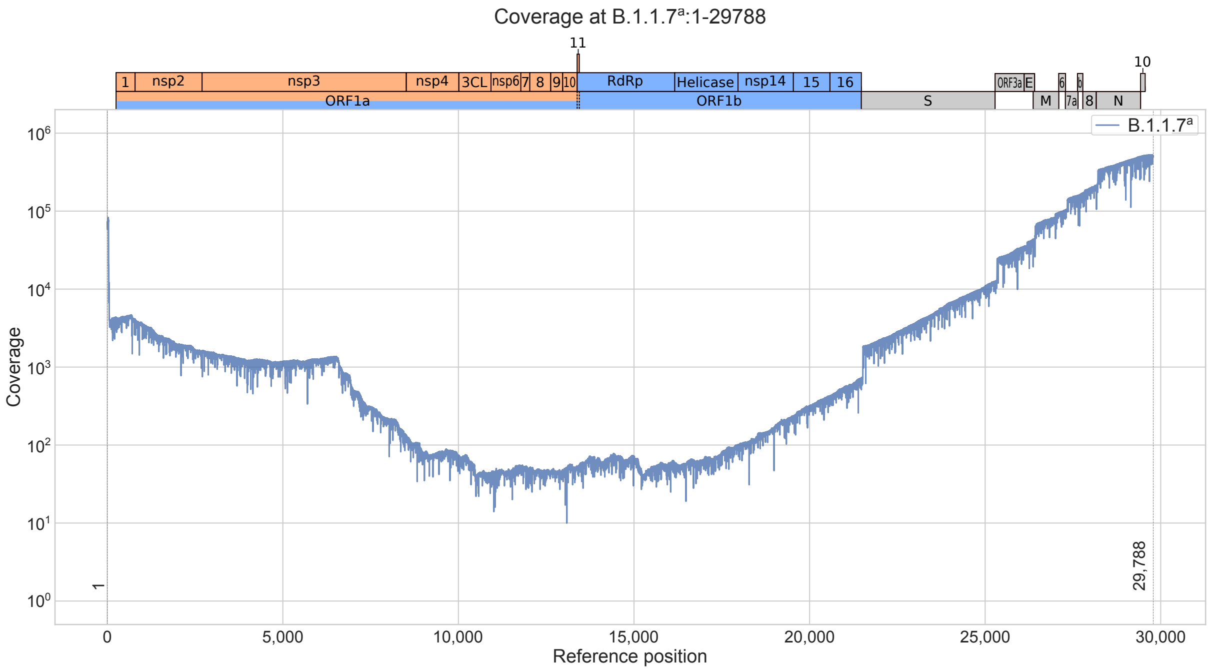


Figure S6: Coverage of B.1.1.7ª Alpha. The coverage is very high near the 5' and 3' ends of the genome and low in the end of ORF1a and start of the ORF1b coding regions. The coverage follows the expected pattern of corona viruses [11, 10]. A low coverage means less data points for the signal distribution approximation. We used a coverage threshold of 10 reads. If the coverage is very low, than the risk increases, that a significant shift is observed by chance. This plot was created using `fastcov 0.1.3` (`https://github.com/RaverJay/fastcov`) and the `mapping.bam` file from the pipeline.

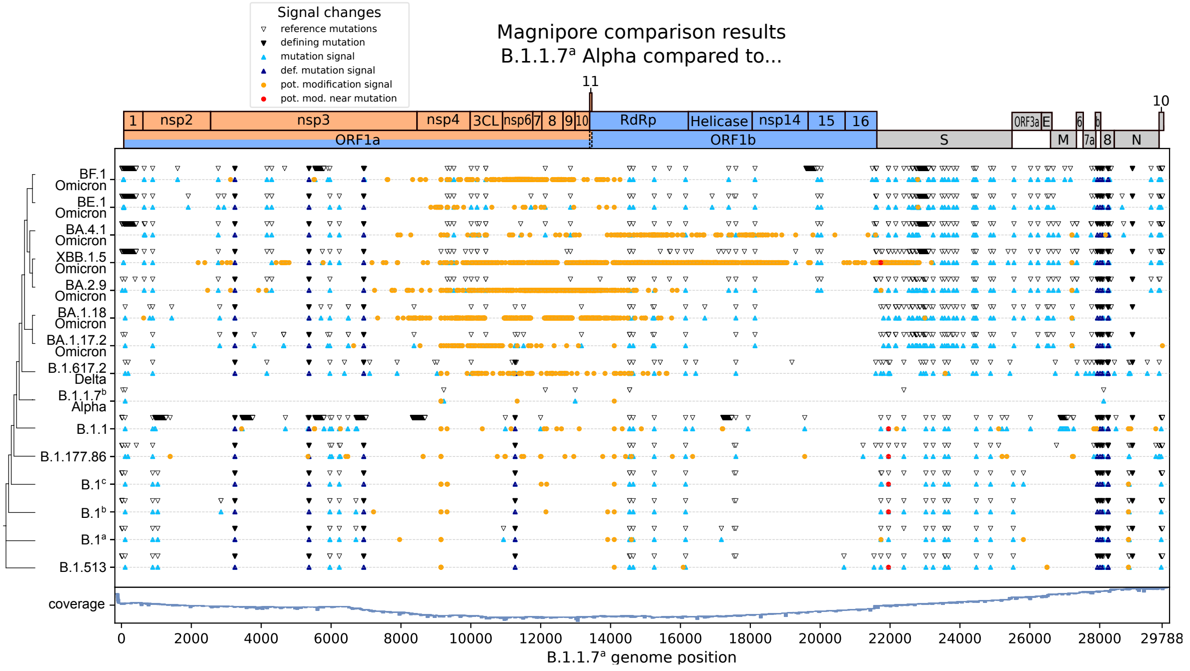


Figure S7: The output of `Magnipore` without a coverage filter yields many potential modification sites in low coverage regions. This indicates, that the approximated signal distributions in low coverage regions are inaccurate or skewed leading to false positive sites between samples.

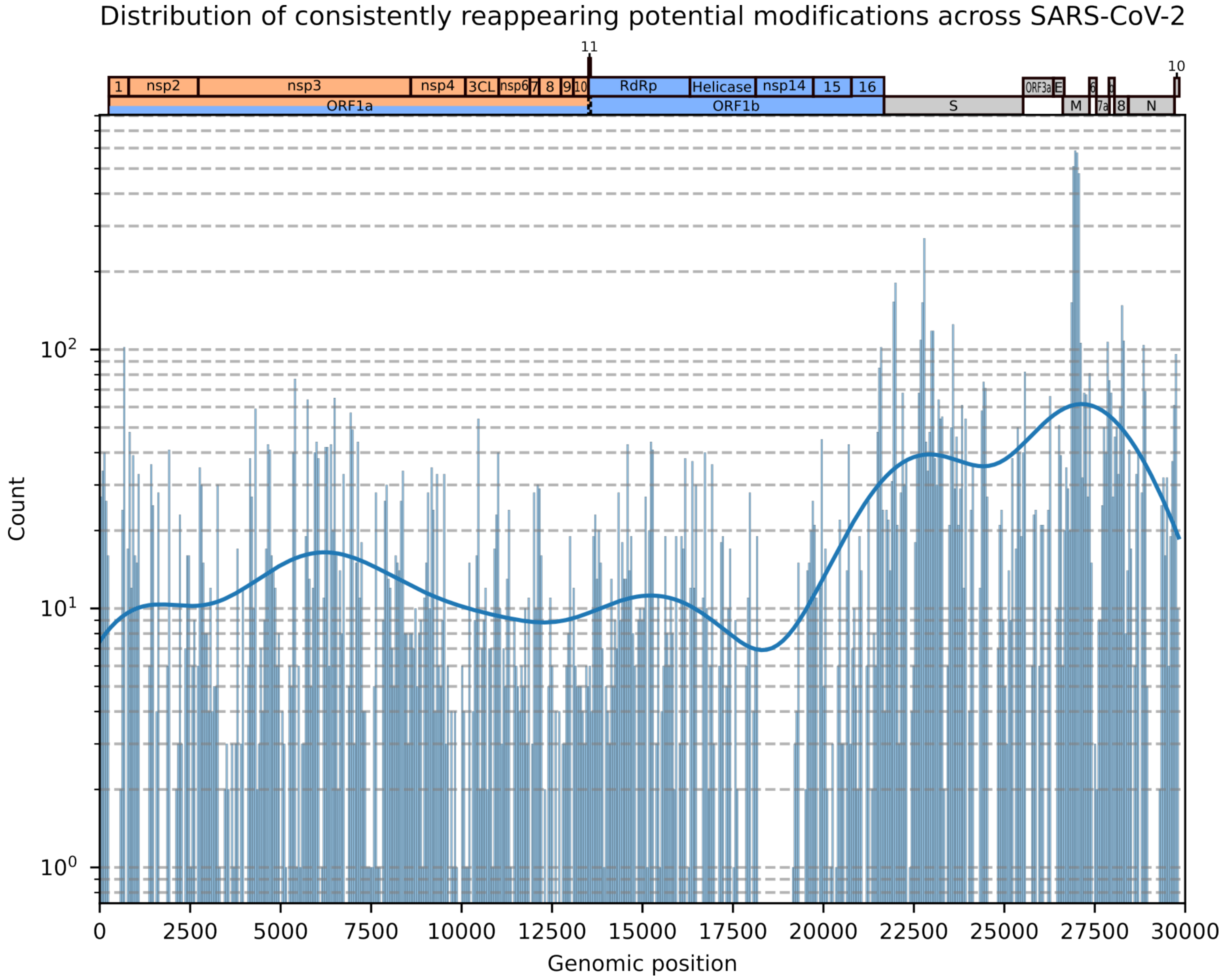


Figure S8: Potential modification sites of all SARS-CoV-2 comparisons in this study. We counted the potentially modified sites for all samples from the potential modification tables in the OSF database and plotted their counts according to their genomic position. Usually the counted genomic position must be corrected for mutations like insertions and deletions between the samples and variants. We did not correct for such mutations. Nevertheless, a high accumulation of potential modifications can be seen near the 3' end of SARS-CoV-2s genome, especially in the S and M gene. Each bar in the plot is a bin of 50 genomic positions.

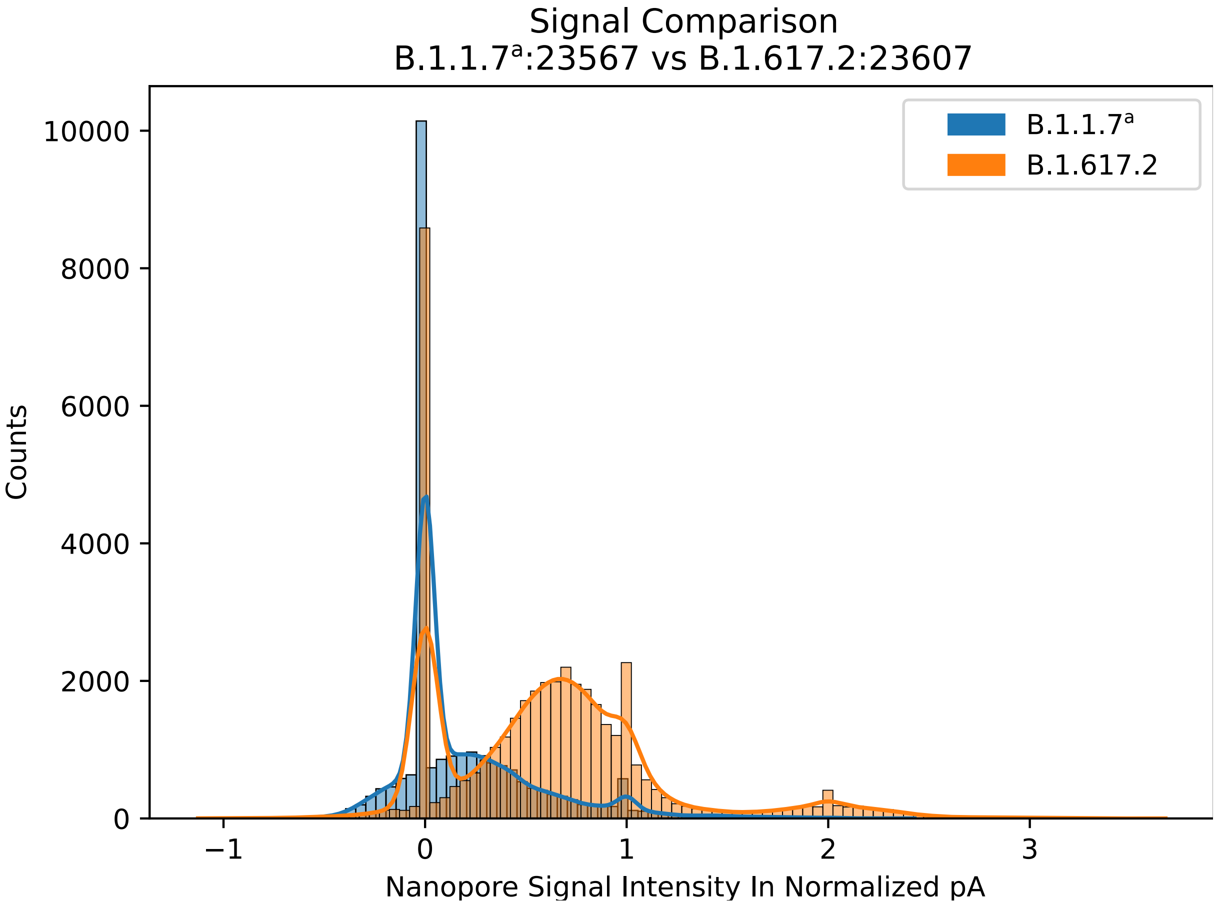


Figure S9: Original signal distribution for position 23 567 in samples B.1.1.7[a] compared to the signal distribution for position 23 607 in B.1.617.2 Delta (ORF: S, Fig 6). `Magnipore` detects a significant site at these aligned positions. It is classified as a potential differential modification as no mutation is present in the genomic context. A significant signal shift between the samples is visible, while both samples show peaks around the normalized pA of 0 and 1. These peaks could result from inaccurate signal segmentation. Additionally, the distributions spread very far and do not behave normal. Another approximation than the Gaussian distribution could be advantageous to describe the signal distribution and analyze such sites.
